# SimTracker tool and code template to design, manage and analyze neural network model simulations in parallel NEURON

**DOI:** 10.1101/081927

**Authors:** Marianne J. Bezaire, Ivan Raikov, kelly Burk, Caren Armstrong, Ivan Soltesz

**Affiliations:** Department of Neurosurgery, Stanford University, Stanford, CA USA 94305.; Department of Anatomy and Neurobiology, University of California, Irvine, CA USA 92697

**Keywords:** computational, modeling, NEURON, hippocampus, GUI, MATLAB, organizing, versioning

## Abstract

Advances in technical computing enable larger and more detailed neural models that can incorporate ever more of the rapidly expanding body of quantitative neuroscience data. In principle, such complex network models that are strongly constrained by experimental data could direct experimental research and provide novel insights into experimental observations. However, as network models grow in complexity and scale, the necessary tasks of development and organization become unwieldy. Further, the models risk becoming inaccessible to experimentalists and other modelers, and their results may then be seen as less relevant to experimental work. To address these obstacles, we developed a tool for managing simulations called SimTracker. It supports users at each step of the modeling process, including execution of large scale parallel models on supercomputers. SimTracker is suitable for users with a range of modeling experience. SimTracker can be a valuable modeling resource that promotes iterative progress between experiment and model.

## Introduction

Rapidly accruing experimental data and expanding computation power are enabling neural network models to grow in complexity and scale (Beltrame and Koslow, 1999; Kandel et al., 2013; Ferguson et al., 2014). This growth makes more difficult the tasks associated with model development and organization of results. Furthermore, models of high complexity risk becoming irreproducible or less accessible to other modelers and experimentalists (McDougal et al., 2016). To address these challenges, here we introduce a novel simulation management tool called SimTracker. SimTracker manages the complexity of developing, executing and organizing results from large scale, detailed simulations executed on multiple supercomputers and addresses the problem of making detailed neural models accessible to experimentalists. Additionally, SimTracker can track hundreds of parameters in these complex models and can organize, analyze, and store results from hundreds or thousands of simulations executed with multiple model configurations. For example, SimTracker has been instrumental in characterizing the interactions between interneurons and principal neurons in a full-scale CA1 network model (see use cases later).

SimTracker works with a flexible network model template written in the NEURON simulation programming language (Carnevale and Hines, 2006), one of the most popular and well-supported programs which has been used in almost 1,700 publications as of September 2015 (Carnevale and Hines, 2015). SimTracker supports the modeler through every step of the modeling process, from designing the model and writing the code to executing the simulations and analyzing the results. SimTracker even helps modelers execute large scale, parallel neural network simulations on supercomputers. Additionally, SimTracker works with code versioning systems to track the code and parameters used to run every simulation, ensuring that each simulation is fully documented and that any model result can be reproduced.

Further, SimTracker provides additional benefits to modelers. Importantly, it addresses the problem of making detailed neural models accessible to experimentalists. SimTracker and its associated model code template allow modelers to quickly characterize their model components in experimental terms, including activation and inactivation curves for ion channels, paired recordings for synaptic connections, and current injection sweeps for intrinsic properties of single cells. In addition, SimTracker is suitable for those with limited modeling experience, providing a structured workflow that can serve as a didactic tool to introduce new modelers to good programming practices and to concepts and pitfalls associated with network models and parallel computing.

## Results

### SimTracker assists with the entire model development process

SimTracker supports the processes associated with model development and execution: design, execution on local and supercomputers, organizing and documenting model configurations and results. It also supports the processes associated with analysis of results: linking results with specific simulations to track data provenance, organizing results, and providing both standard and custom analyses for each simulation. From within SimTracker, a user can specify the parameters to use in the simulation, execute the simulation, load its results, and produce standard figures to analyze the network behavior (Figure 3). SimTracker saves a record of every simulation, including those that have been designed but not yet executed.

**Figure 3:**
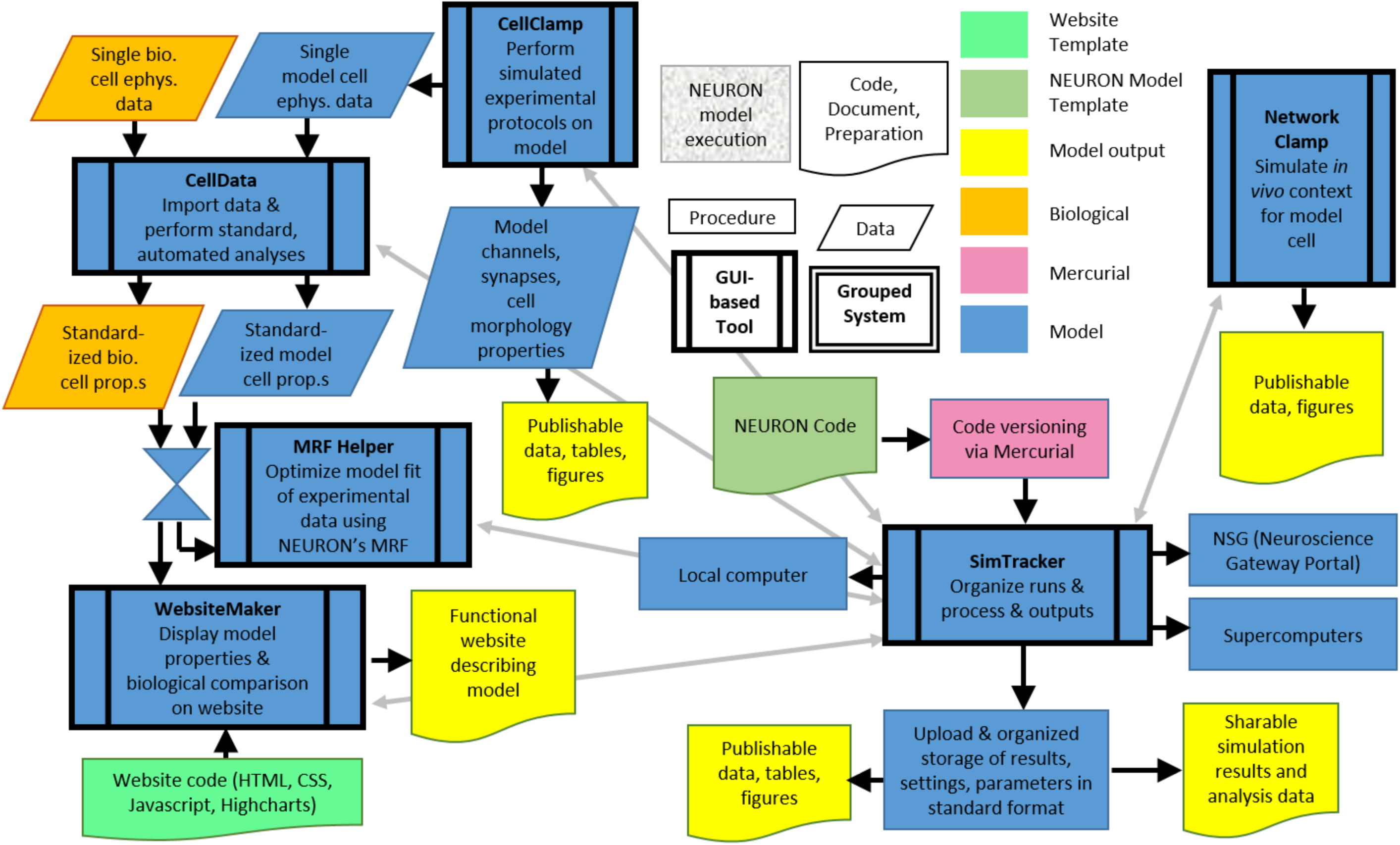
A brief workflow illustrates how SimTracker, its various components, and the NEURON code all work together to support the complete model development process.

For parallel network runs that require a large supercomputer for execution, SimTracker generates and submits the job script to the supercomputer’s batch queuing system. When the simulation has completed, SimTracker can download all the results from the supercomputer. SimTracker can also be used with the Neuroscience Gateway portal (NSG, https://www.nsgportal.org/); it will package the simulation directory in the format necessary for submission to the NSG and can extract the results from an NSG output package.

SimTracker works with Mercurial, an established code versioning system, to track the exact code used to execute each simulation. It uses configuration sets to specify cell numbers, connectivity, and synapse kinetics of the model network. The modeler can edit these configuration sets as well as all parameters used in the simulation (Figure 4). These two abilities together allow complete documentation and replication of any simulation. SimTracker also ensures that the code version on the supercomputer where code will run matches the version on the user’s personal computer, and that the parameters to use for the run are specified on the supercomputer and that the necessary configuration sets are present.

**Figure 4:**
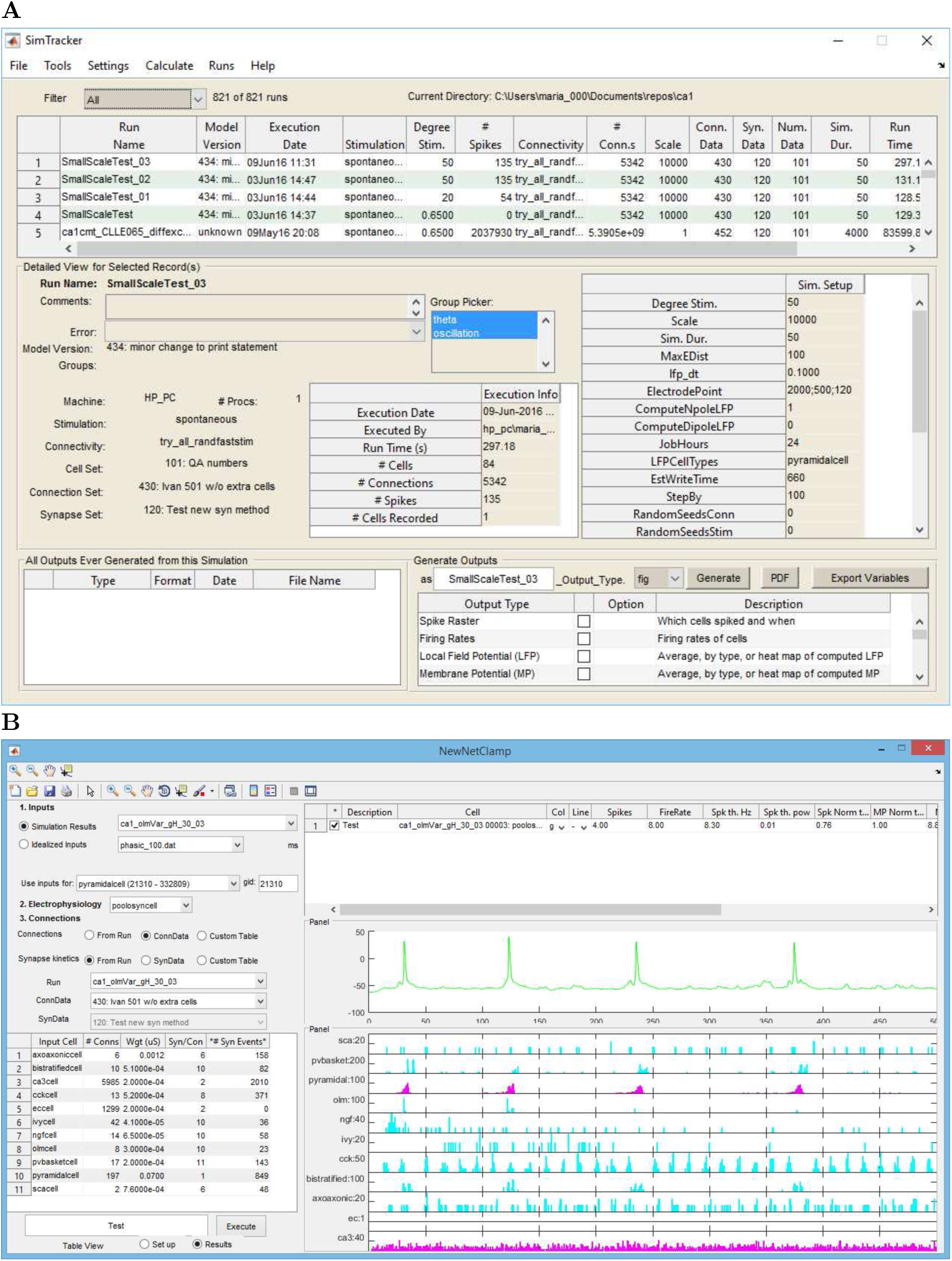
SimTracker and Network Clamp. (A) The main SimTracker interface, along with its parameter menus and tools used to specify the parameter datasets for use in the model. (B) the Network Clamp tool for running Network Clamp simulations.

SimTracker can produce a variety of outputs for each simulation, including spike rasters, connectivity matrices, or single cell traces. Additionally, users can add their own custom scripts to analyze their simulation results. Each time that SimTracker produces a figure from a simulation, it logs which simulation produced the figure, the name of the figure file, and its location. Logging the details of figure generation enables better tracking of figure origins that becomes important once many simulation results are produced. SimTracker assists the user with other aspects of organization as well. It can archive, reload, and back up simulation results. Users can easily share the results of simulations with each other using SimTracker’s import and export functions. Simulation results can also be uploaded to the web so that others can analyze them without having to rerun the whole simulation or access a supercomputer themselves.

Within SimTracker (Figure 4A) are four additional tools to support the model development process: NetworkClamp (Figure 4B), CellData (Figure 5A), CellClamp (Figure 5B), and WebsiteMaker (Figure 5C). CellClamp allows the corresponding analyses to be performed on the model cells. In addition to performing single cell recordings to characterize intrinsic properties of model cells, CellClamp also conducts paired cell recordings to characterize synapses and and recordings of ionic membrane conductances to produce the current/voltage curves and activation/inactivation curves commonly used to characterize ion channels. CellClamp can also prepare a NEURON session for tuning a model cell using NEURON’s Multiple Run Fitter (MRF) tool. The MRF is a powerful tool useful for tuning model parameters to match experimental data, but it can be intimidating to configure. Therefore, SimTracker can set up a MRF session with user-specified parameters, gently introducing the modeler to the process of model optimization using MRF.

**Figure 5:**
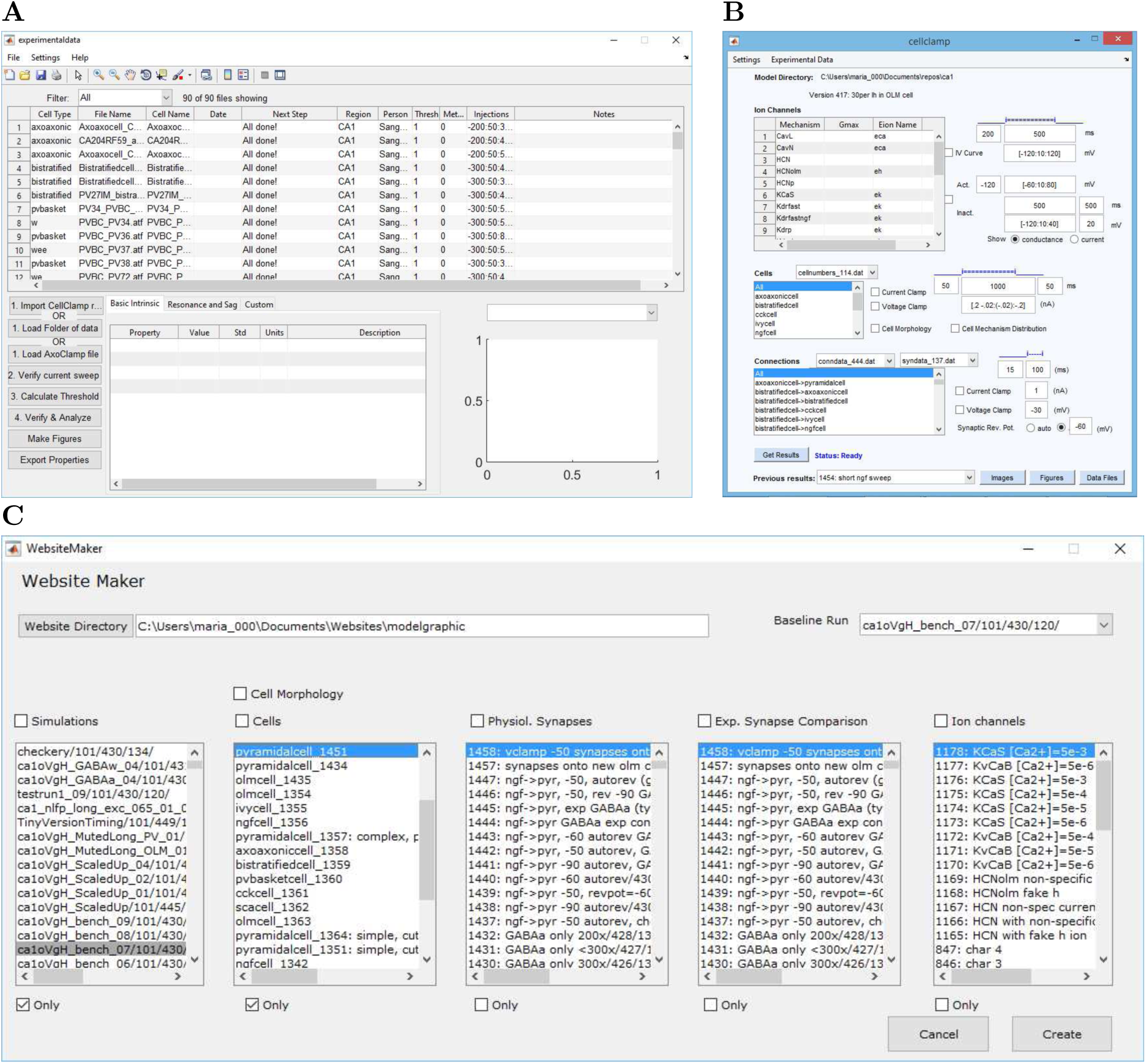
Additional tools within SimTracker. (A) Cell Data tool for analyzing intrinsic electro-physiological properties using raw AxoClamp files of the data from single cell current injection sweeps. (B) Cell Clamp tool for characterizing cells, channels, and synapses by subjecting them to biological experimental protocols. (C) Website Maker tool for generating a website that displays the model characteristics and compares model and experimental network components.

### Network Clamp

NetworkClamp enables a single cell to be ‘Network Clamped’ to the inputs of a full size network. In brief, the NetworkClamp tool allows the user to specify a cell of interest, which network inputs to give the cell, and the spiking activity patterns of the network inputs (whether idealized inputs, actual spike trains from a simulation or experiment, or some other pattern). A deailed example of Nework Clamp usage is presened below.

#### Interneuronal contributions to theta oscillations in simplified models derived from the full-scale virtual CA1 network

Network Clamp allows a modeler to extract a single cell from the full-scale network, called the ‘cell of interest’, while maintaining all its spatially distributed synaptic inputs and their cell type-specific presynaptic activity patterns, so the cell of interest remained within the context of the network. To conserve network context, Network Clamp can maintain the same number and properties of input synapses to the cell, including the synaptic diversity for different presynaptic cell types (kinetics, amplitude, location, reversal potential). However, Network Clamp also has the ability to receive an arbitrar network input to recreate other contexts besides that of the model network.

In order to gain additional insights into the role of interneurons in the generation of the hippocampal theta rhythm, we took advantage of our uniquely data-driven, full-scale CA1 model (Bezaire et al., 2016) by rationally deriving simpler models from it. Our general aim was to develop a new modeling approach that would yield simpler models with dramatically fewer parameters and minimal computational resource requirements, without losing the strong biological realism of our data-driven full-scale model. For this new approach, we extracted a single cell from the full-scale network, called the ‘cell of interest’, while maintaining all its spatially distributed synaptic inputs and their cell type-specific presynaptic activity patterns, so the cell of interest remained within the context of the network. Specifically, in order to conserve network context, we maintained the same number and properties of input synapses to the cell, including the synaptic diversity for different presynaptic cell types (kinetics, amplitude, location, reversal potential). We also replaced the presynaptic interneurons that activate those synapses with realistic spike trains whose properties were based on experimentally observed spike trains recorded from identified interneurons during theta (Figure 1; see Methods). Importantly, the excitatory synaptic inputs to the extracted CA1 pyramidal cell still received Poisson-distributed, arrhythmic presynaptic spike trains, as in the full-scale model. This strategy ensured that the model cell received the same number and spatio-temporal character of inputs as it would in the full network. We termed this modeling method ‘Network Clamp’, because the activity in the rest of the network was essentially dynamically clamped, with changes in the spiking output of the extracted pyramidal cell not having an effect on the incoming signals from the rest of the network. The key advantage of the approach was that the extraction of a single pyramidal cell from the CA1 network drastically reduced the number of free model parameters, while retaining most of the numerous experimental constraints about CA1 pyramidal cell single cell anatomy, connectivity and electrophysiology.

**Figure 1:**
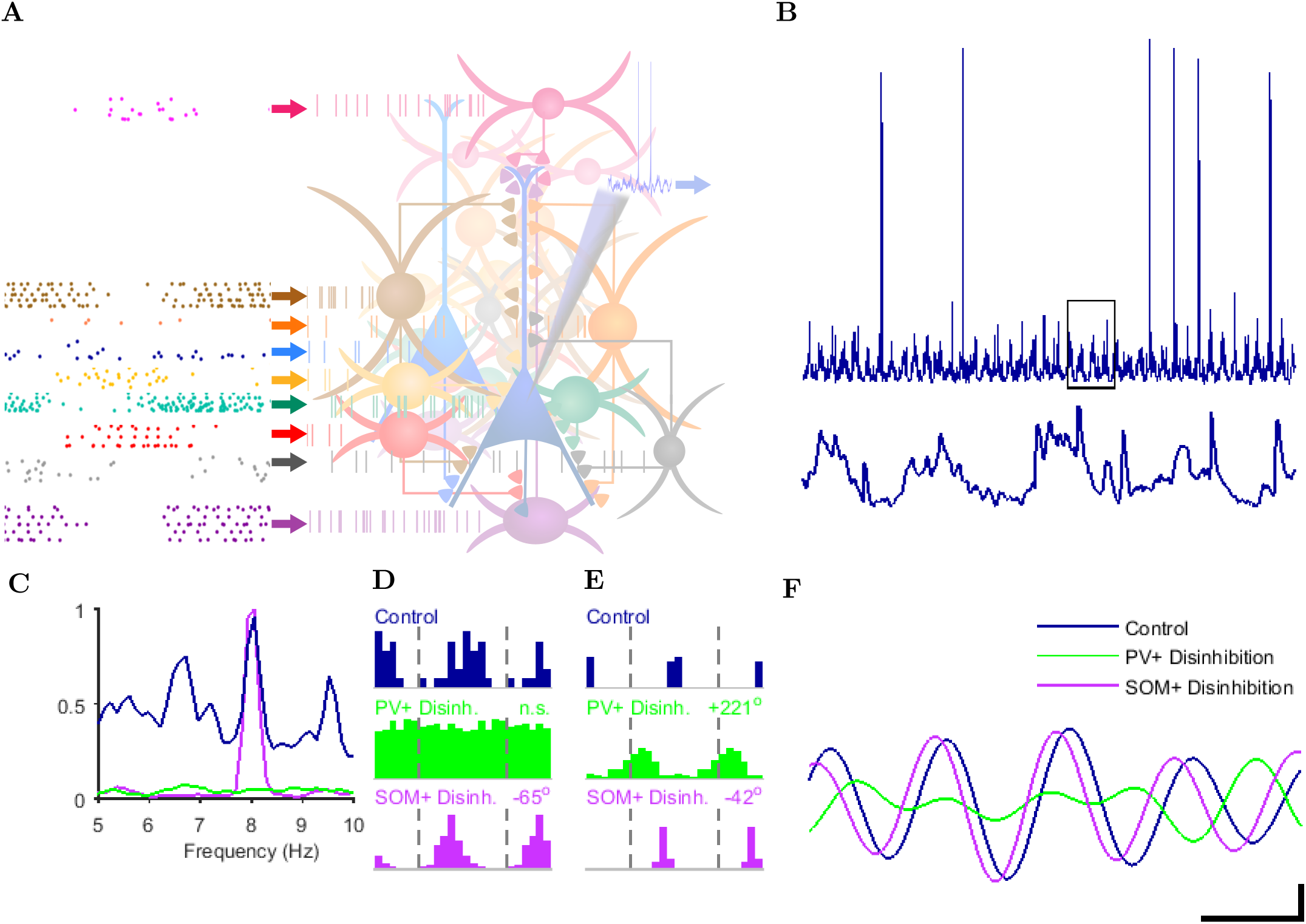
Network Clamp. (A) Conceptual diagram of the network clamp shows a single cell as the focus, with biologically-constrained convergence onto the cell as if it were part of the full network, and with the presynaptic connections to the cell firing realistic spike trains as if embedded in the network. The intracellular somatic membrane potential of the cell of interest is recorded. (B) Theta oscillations are seen in the membrane potential of the cell within the context of a full network during theta (top), while the inset (bottom) shows simultaneous high frequency oscillations. Scale bar in F represents 1.04 s and 9.4 mV for top trace in B and 100 ms and 2 mV for bottom trace in B. (C) Normalized Welch’s Periodogram of the spike density function of the pyramidal cell of interest shows that the PV+ basket cell disinhibition causes a loss of theta rhythmic firing while the O-LM cell disinhibition does not (with other cell inputs remaining the same). (D) Theta phase firing probability in control and disinhibition states shows greater phase shift and loss of phase modulation due to PV+ basket cell disinhibition compared to O-LM cell disinhibition. Note that the spike phase were calculated relative to the intracellular membrane potential oscillation here in contrast to the calculations for full network results, which were calculated relative to the extracellular LFP. (E) To ensure the observed cell type-specific differences in the effects of reduction of interneuronal inputs on theta oscillations were not simply an artifact of the increased firing itself, we blocked the sodium channels in our Network Clamped pyramidal cell and analyzed peak depolarization times instead of spike times. Again, we found a markedly stronger effect following PV+ compared to SOM+ disinhibition in both the phase shift of peak membrane potential (analogous to spike time) and in the drop in theta modulation level. Histograms display the theta phases of the peak depolarizations of the intracellular somatic membrane potential. (F) Filtered membrane potential trace, as well as filtered trace of the potential with 90% PV+ basket cell inhibition removed (green) or 90% O-LM cell inhibition removed (purple) contrasts the different phase shifts caused by each disinhibition. For traces in panel E, scale bar represents 100 ms and 1 mV.

## CellData

CellData reads in Axo-Clamp data files from experimental cell recordings and analyzes standard electrophysiological properties to provide constraints for model cell development. The CellData tool can be used independently from SimTracker by experimentalists who only wish to automate their cell analysis workflow. However, it also integrates with the other components of SimTracker, for example by providing constraints for a MRF optimization session or for supplying comparisons and biological cell behavior to the website template to showcase the fitness of the network model components.

## WebsiteMaker

Finally, WebsiteMaker allows one to populate the contents of the datasets used in a specific network simulation and the results of CellClamp characterizations into a website template, generating the code for an attractive and interactive graphical online tour of model network components such as seen at http://mariannebezaire.com/models/ca1/. The website is intended to provide model transparency and give readers a good sense of the inner workings of the model.

## SimTracker is flexible and can support a variety of projects

To highlight the broad utility of SimTracker for models simple and complex, large and small, we describe two disparate modeling use cases here. First, SimTracker can be used as an introduction for new modelers by following tutorials 1-5 in the Tutorials section below. Through these tutorials, users will become familiar with basic concepts of modeling, such as designing the model, implementing it, testing it, running the code in parallel for larger networks, and analyzing the results. Users will also learn about organization of the model, code versioning, and tracking of parameters.

Tutorials 1-5 use the same ring network model that Hines and Carnevale (2008) employed to introduce strategies for parallelizing NEURON code. In this work, we show how our model NEURON code template can be used with SimTracker to design and execute the same ring demo network simulation, producing the same results as those obtained in Hines and Carnevale (2008).

SimTracker can also be used to support entire projects involving modeling a complex network at full scale with numerous biological constraints (see tutorials 8-10 regarding large scale simulations on supercomputers and tutorials 11-14 tegarding incorporating parameters and constraints). To illustrate the capabilities of SimTracker regarding such large and detailed models, we will briefly describe how we used SimTracker in our recent work addressing spontaneous theta oscillations in a model of the isolated hippocampal CA1 network (Bezaire et al., 2016). The first question we explored was whether the constrained, biologically detailed, full-scale CA1 model was capable of spontaneous physiological oscillations. We varied the non-rhythmic, tonic excitation to the network and found a range of excitation levels where the network developed a stable, spontaneous theta rhythm. Critical to this process were SimTracker’s capabilities for tracking model configurations. We used SimTracker to track the many simulations we ran as we investigated the effects of various excitation levels and the effects of using different values for those biophysical parameters for which there was little or no available information. After completing our simulations, we decided to add code to calculate the local field potential (LFP) generated by our model network. Because SimTracker tracks the provenance of each simulation thoroughly, we were able to quickly rerun all our simulations with the LFP code added to record the LFP trace associated with each simulation. Furthermore, SimTracker allowed us to efficiently apply our custom analyses to each simulation, so that we could quickly obtain the spectral analysis of the simulated LFP signal and of the pyramidal cell spike density function to determine how the network dynamics depended on various network components. SimTracker’s standardized visualization and calculation capabilities were instrumental in plotting and comparing the results of these two distinct methods in order to confirm that biophysical oscillations were present in the model.

We also investigated in Bezaire et al. (2016) whether the distinct interneuronal types were necessary for biophysical oscillations to develop in the network. We constructed a number of variants of the basic network model, gradually decreasing the diversity of interneuronal species in the network, and conducted a number of simulation runs at varying excitation levels. As a result, we established that interneuronal diversity is a prerequisite for stable theta oscillations. As this study necessitated a large number of similar network configurations, as well as a large number of simulation runs for each configuration, SimTracker was crucial to keeping track of all network configurations, and corresponding simulation runs and results.

During our initial model development and validation, we used SimTracker to characterize model components as shown in Figure 6. When it came time to share our model results, SimTracker was able to load the model component characterizations into a website template, creating a resource for anyone who wishes to understand our model network better.

**Figure 6:**
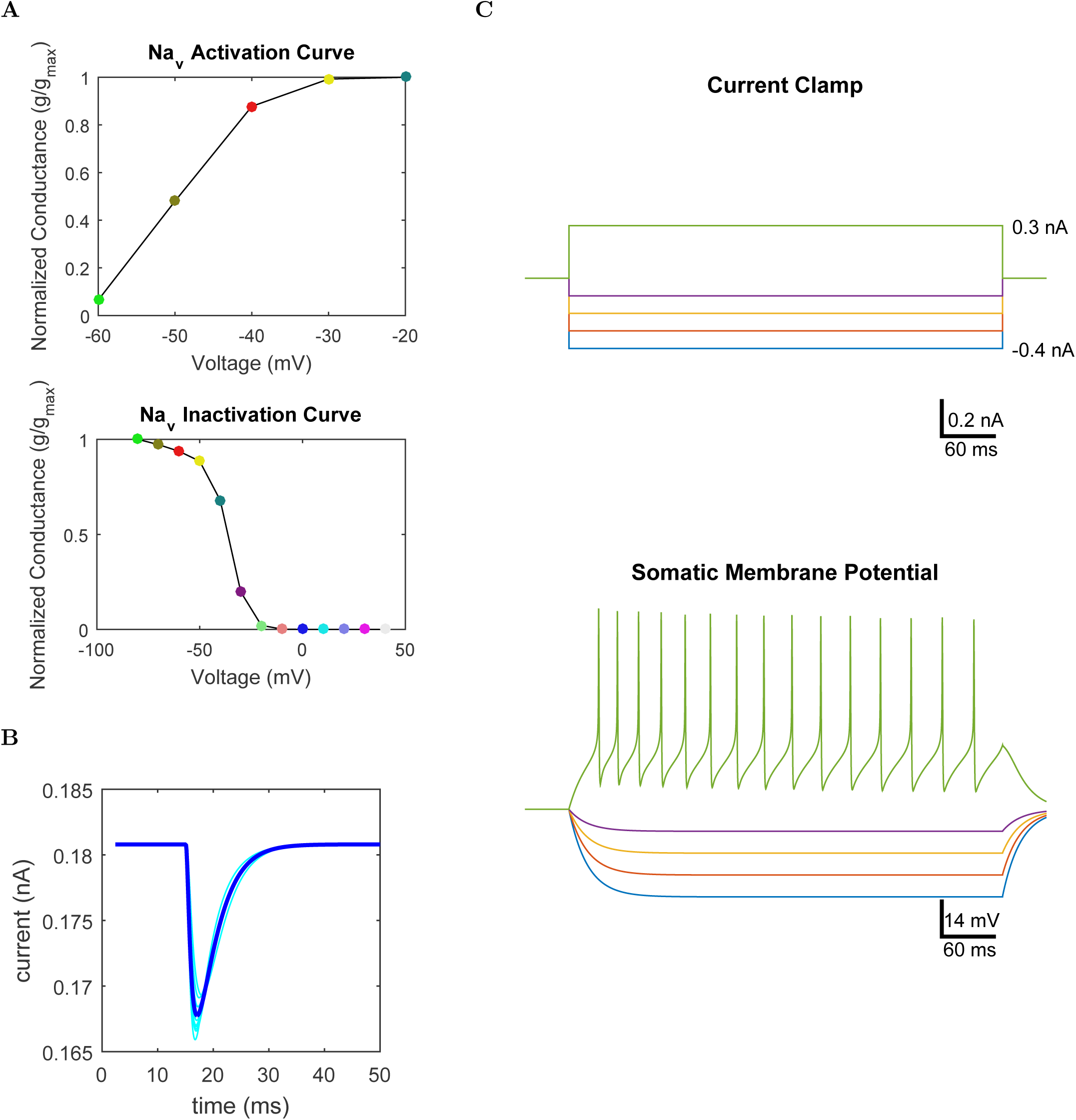
Various aspects of the model network can be characterized in experimental terms.(A) Activation (top) and inactivation (bottom) curves for a voltage-gated sodium channel. (B) Average and individual postsynaptic currents (n=10) for connection from an excitatory afferent to a pyramidal cell. (C) Somatic current injection sweep for a model neurogliaform cell.

In addition, the CellData tool within SimTracker can be used by independently by experimentalists with no modeling experience to partially automate the analysis of AxoClamp recordings from single cell current injection sweeps. CellData reads in AxoClamp files and measures many intrinsic properties using standard calculations, as detailed below in section ‘The CellData tool within SimTracker can be used independently by experimentalists’ and also as illustrated in Tutorial 6. For modelers, the CellData can integrate with the other SimTracker tools to streamline the process of tuning model cells to experimental constraints.

## SimTracker NEURON code template is well parallelized and set up for optimization on supercomputers

An important resource associated with SimTracker is the NEURON code template itself. The NEURON code template and SimTracker tool were developed together to streamline the entire modeling workflow and minimize the possibility for errors in simulation tracking and design. However, the NEURON code template can be used independently of SimTracker for modelers who only need sample code upon which to base their project.

The NEURON model code template is well parallelized for networks containing many more cells than the number of processors used to execute the parallel simulation. We tested the model’s performance by running the full scale CA1 network model on differing numbers of processors (Bezaire et al., 2016). We found an almost linear relation between the number of processors used and the wall-clock time required for the simulation to complete. Stated another way, the total number of processor-hours (hours of computation performed by each processor used in the simulation) remained almost constant and independent of the number of processors used. This linear relation indicates that the model code was only directing processors to perform necessary computations and that there was little overhead associated with including additional processors in the computational job. Only with the largest job size of 3,488 processors did we see a drop in efficiency, possibly due to the increased communication time required with more processors. As seen in Figure 7, the relation between processor number and wall-clock time for the three smallest job configurations can be fit to Equation 1 with r^2^ = 1.00:

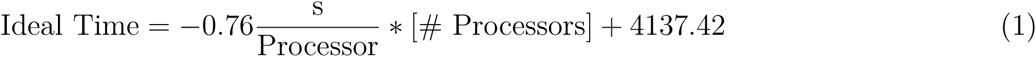

**Figure 7:**
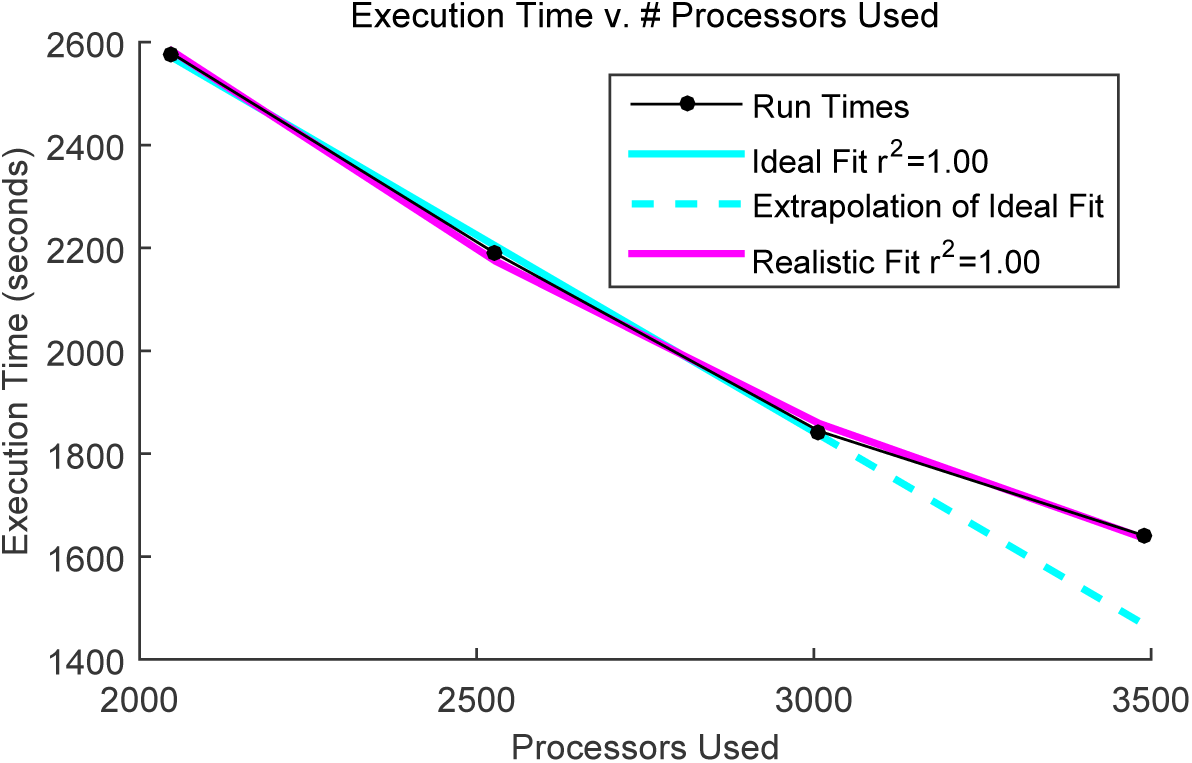
There is a near linear relation between the execution time and the number of processors used for a parallel simulation of our large-scale network model code, indicating well-parallelized code. As the number of processors increases further, there is an overhead associated with exchanging information between additional processors that causes a decrease in execution efficiency. The ideal fit was computed from a linear equation, while the realistic fit was computed from a second-order polynomial. This figure includes results from 4 simulations using unique numbers of parallel processors; therefore, n=1 for each condition. We present the raw data (time required for simulation) for each condition and then fit both linear and polynomial lines to the data, computing Pearson’s coefficient to measure the fits.

However, the gains made from adding more processors fall off for the largest number of processors, so including all job sizes, a quadratic equation fits better as seen in Equation 2 with r^2^ = 1.00:

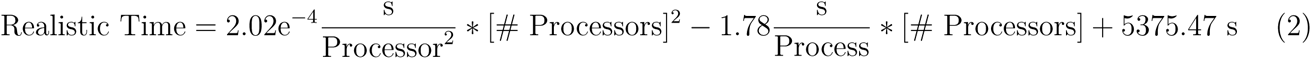

To ensure good parallelization, we also calculated the load balance of the simulation jobs to ensure that all processors performed a similar number of computations throughout the course of the simulation. Indeed, we found that the load balance of the simulations exceeded 99.8% on each of our benchmark runs. These strong results were expected because the number of cells in our model was far greater than the number of processors on which the simulation was ran, enabling us to evenly distribute the cells among the processors to achieve equivalent computational demands on each processor.

Finally, we also measured the time required for processors to communicate with each other. Processors must communicate with each other when a postsynaptic cell and a presynaptic cell are not on the same processor. In that case, the spike time of the presynaptic cell must be sent to all processors that own any of its postsynaptic cells; therefore this communication time is also called the spike exchange time. With a well parallelized model, it may be attractive to increase the number of processors in a job to the limit of the supercomputer with the hope of steadily decreasing the wall-clock time needed to complete the job. However, there is theoretically a slight cost of time to adding more processors because more spike exchanges will then be necessary, and these spike exchanges take time. In practice, we found that the exchange time was always less than 2 seconds regardless of whether we used 1,728 processors, 3,488 processors, or something in between, and there was no significant correlation between exchange time and number of processors for the job configurations we tested.

We also ran the code on a variety of computers, and we tested accessing the supercomputers directly or via the Neuroscience Gateway. We found the code worked well on each computer that we tested. We specifically ran the CA1 network model on Stampede and Comet and found both computers to be equally suitable. Accessing the computers via the NSG did not change the model performance or results. The same model results were obtained on Stampede and Comet. See Figure 2. In general, a simulation lasting 100 ms in the full size CA1 network model required about 1,500 processor hours (or about 40 minutes using 3,008 processors), and needed 2.8 TB of RAM.

**Figure 2:**
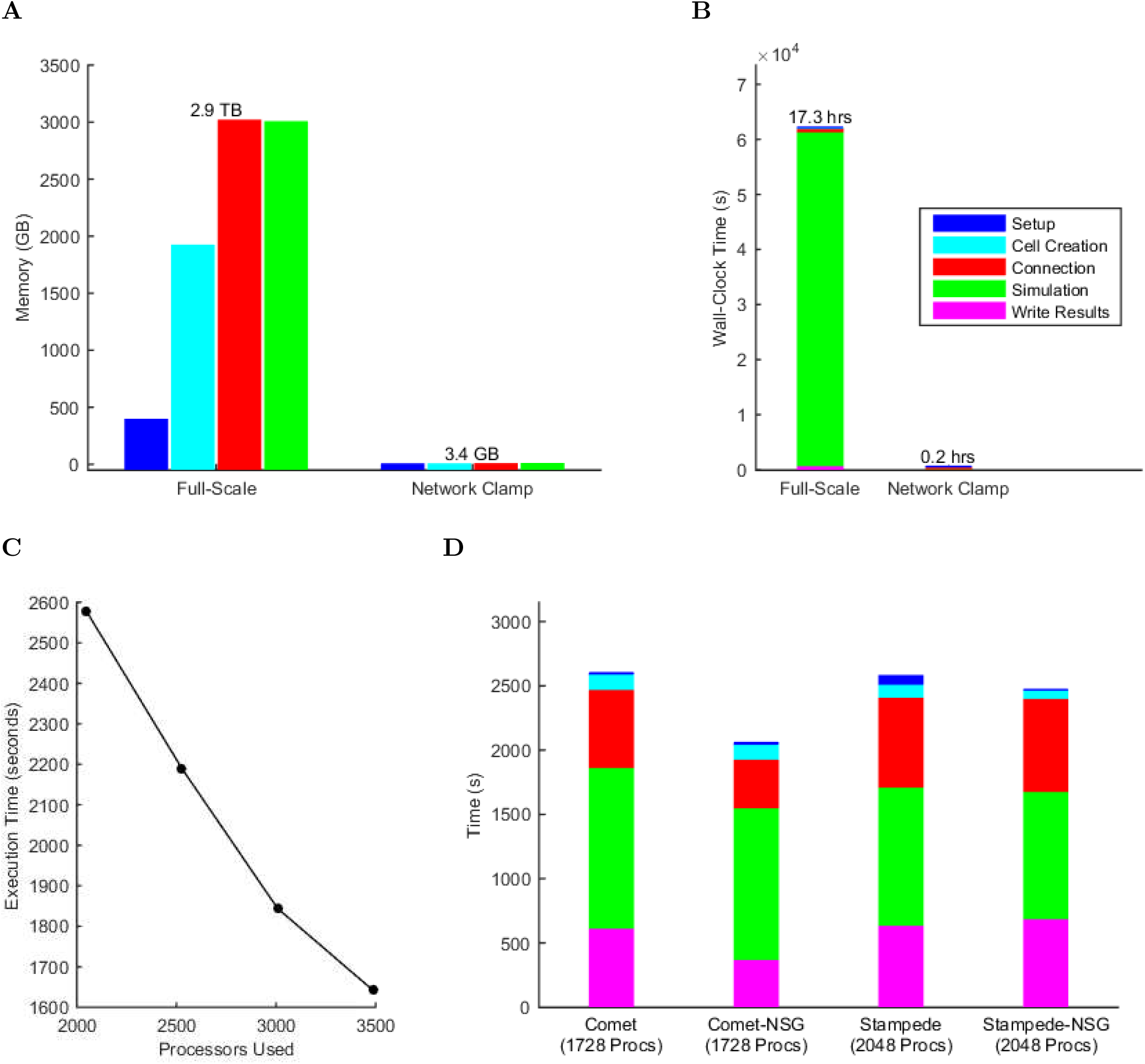
PModel Performance. (A-B) Comparison of full-scale and Network Clamp requirements for (A) memory and (B) wall-clock time required for computation. (C) Wall-clock time requirements of the full-scale model when run on various numbers of processors (100 ms simulation). (D) Running the same 100 ms simulation on different supercomputers (NSG = Neuroscience Gateway portal, Procs = processors). These times are for simulations that performed a simplified LFP calculation; LFP calculation used in the model analyses tracked all local pyramidal cell compartments and required additional run time.

We took many steps to decrease the time and memory requirements of our model, in addition to parallelizing the code. For example, we compiled the steps of the network creation that were performed repetitively: assigning each cell to a location in space, and determining which cells to connect while respecting type-specific connection preferences and axonal distributions and extents. We also made the simulation more efficient by using cache efficiency and, for simulations that meet certain requirements, spike or cell ID compression.

Because different networks will exhibit different dynamics and require different resources, we have written the code so that it is a simple task to characterize the network spiking activity and then reconfigure the spike data transmission to be most efficient for that particular network.

## The CellData tool within SimTracker can be used independently by experimentalists

Not only does the CellData tool interface with CellClamp and the MRFHelper to assist with tuning model cells, but it also functions independently as a tool to partially automate analysis of AxoClamp files from single cell current injection sweeps. Experimentalists can benefit from the CellData tool even if they are not planning to use any modeling in their research.

The CellData tool is designed to be used in conjunction with experimental protocols in which single cells are whole-cell patch clamped, subjected to current injections at a range of levels to characterize their hyperpolarized, subthreshold, and suprathreshold responses, and recorded using AxoClamp software. CellData has been tested by previous experimentalist end-users to calculate intrinsic excitability data for a large number of cells (Armstrong et al., 2015). CellData reads in AxoClamp files in ABF or ATF format, parses the current injection information and the recorded membrane potential information, and then analyzes the behavior of the cell at each current injection level. Users can select from a variety of methods for automatically finding the action potential threshold, with an opportunity to review all automatically selected threshold points (as well as sag, depolarization, AHP (after-hyperpolarization) and action potential peaks or troughs). Upon confirming that CellData has accurately interpreted the recorded membrane potential traces, users then receive a table of the cell’s intrinsic properties and the option to produce graphs of the cell’s behavior. Multiple cells can be processed and displayed at a time, and the calculations used to produce the intrinsic property table are clearly documented and standardized. The table can be exported from CellData in a variety of formats, including ones compatible with Microsoft Excel, Mac OS Pages, and tab-delimited or CSV formats, while the figures can be exported in all common figure formats or saved as MATLAB figure files. See tutorial 6 for more information.

## SimTracker is available and accessible to all

The MATLAB code for running SimTracker is publicly available from https://bitbucket.org/mbezaire/simtrackercode. In addition, compiled versions of SimTracker can be found on http://mariannebezaire.com/simtracker/ with a variety of additional resources, and links to SimTracker are available on SimToolDB (a known database for simulation tools) at https://senselab.med.yale.edu/SimToolDB/showTool.cshtml?Tool=153281, MATLAB Central (link under review), and other websites. SimTracker is available as MATLAB source code or as a compiled, stand-alone application for those without MATLAB licenses. SimTracker conforms to the Open Source Definition and is licensed under the MIT License, which has been approved by the Open Source Initiative (https://opensource.org/about). Every effort has been made to document SimTracker usage for users with a variety of technical abilities. In this article, we will introduce the functionality of SimTracker with a series of tutorials that are sufficient to foster proficiency in an average neuroscientist. For more support, potential users can consult the detailed instruction manual and other resources online at http://mariannebezaire.com/simtracker/.

SimTracker currently works with the NEURON simulator (Carnevale and Hines, 2006) and interfaces with several other applications throughout the development workflow of a simulation, including the batch queuing software of several supercomputers, Mercurial, Microsoft Excel, and the Multiple Run Fitter tool within NEURON. It could be customized to work with other simulator software and interfaces with other programs throughout the modeling process workflow.

Though SimTracker is MATLAB-based, users do not need MATLAB to use SimTracker. Compiled, stand-alone versions of SimTracker exist for Windows (32-bit and 64-bit), Mac, and Linux operating systems. In addition, the MATLAB code used to create SimTracker can be freely customized under the terms of the license. SimTracker has been tested by several users on Windows and Linux, using both the compiled and the uncompiled versions. Additional issues found within SimTracker can be reported online within the NEURON forum located at https://www.neuron.yale.edu/phpBB/viewforum.php?f=42.

To streamline the installation of SimTracker, we have created a Docker image (software bundle) for Linux operating systems. This bundle allows the user to install SimTracker, Mercurial, and NEURON all at once, vastly simplifying the setup process. Bundled software packages will soon be available for Windows and Mac OS X operating systems.

## Discussion

One of the main goals of detailed, biologically constrained neural network modeling is to test mechanistic hypotheses generated from experiments and to make new, testable predictions that can guide future experimental work. As our hypotheses about brain function increase in nuance, scope and quantification, computational models can more strongly drive and interpret experimental research. For the interaction between model and experiment to be most effective and consistent, we must be confident in how well our model represents the experimental system and confident that our model implementation matches what we meant to design. We must also recognize the utility of representing the output of our model in experimental terms. SimTracker aids with all of these requirements to ensure tighter coupling between experiment and model.

As quantitative experimental data becomes more prevalent and accessible, modeling at a detailed level (individual cells, detailed synapse and ion channel characterization) is also becoming more common. There are now over 1,679 published neural models implemented in NEURON alone (Carnevale and Hines, 2015), many of them including physiologically detailed cells that incorporate ion channel mechanisms (with characteristic activation, inactivation processes and dependence on voltage or other conditions). Many of these detailed models are of networks, even large-scale networks incorporating incorporate detailed connectivity information (synapse strengths and kinetics) and requiring parallel NEURON and supercomputers.

To track the parameters of these detailed models, as well as to accurately describe and characterize the model in a manner that allows the model to be useful to others and the results to be placed in proper context, model transparency is necessary. Models generally can and should be made more transparent (Merali, 2010), as increased transparency aids comprehension, validation, reproducibility – at the level of the code and at the level of model behavior. However, characterizing models can be time-consuming and even when it is completed, may be done in non-standard ways. Here, we have provided a tool that allows for standard characterization of model components at several levels, from the code itself to the instantiated network components. SimTracker enables all network components to be characterized using computational analogs of common experimental procedures, which serves the purpose of increasing model transparency and also making the model more accessible to non-computational scientists who may be interested in the implications and predictions of the model. This standardized characterization process is also made possible by the organization of the NEURON code into a common template that can be used across a variety of network models. SimTracker can serve as a training device, walking new modelers through the steps required to develop rigorous model code, and introducing them to important concepts and practices they may not have been familiar with. In addition, SimTracker provides standard methods for analyzing model behavior and generating figures for publication.

There are other simulation management resources available, some of which have overlapping functionality with SimTracker, including NeuroConstruct (Gleeson et al., 2007), Net-PyNE (http://www.neurosimlab.org/netpyne/), Lancet (Stevens et al., 2013), NeuroTools (https://github.com/NeuralEnsemble/NeuroTools), NeuronVisio (Mattioni et al., 2012), and Sumatra (Davison, 2012). However, a combination of properties renders SimTracker unique and a necessary addition to the other available tools. SimTracker provides a structured, all-encompassing resource that tackles all aspects of running parallel NEURON network models while still allowing the user relative flexibility in the implementation of the NEURON code through a separate NEURON code template. SimTracker also has features that may increase its accessibility for non-programmers. It is GUI-based (a graphical user interface, such as most applications used by non-programmers) and geared toward users with varying levels of coding experience, while many of the other options are library-based, intended for users who prefer to write their own scripts to make use of the features. The structure and level of support available in SimTracker is made possible by its focused scope. The use of SimTracker does not preclude the use of additional modeling resources: SimTracker has been specifically designed to work with NEURON and its tools, and models created using SimTracker can easily be uploaded to ModelDB (https://senselab.med.yale.edu/modeldb/) or OpenSourceBrain (http://opensourcebrain.org), or ported to NeuroML (Gleeson et al., 2010) and used in NeuroConstruct (Gleeson et al., 2007), as well as ran using the Neuroscience Gateway portal (NSG, https://www.nsgportal.org/). In fact, our NEURON model code templates are currently being implemented in NeuroML by the NeuroML team, after which they will be made freely available on OpenSourceBrain (http://opensourcebrain.org/projects/nc_ca1). The NeuroML version of the code can be immediately used with many of the other simulation helpers listed above for modelers who would like to run the model but already have a preferred workflow for their modeling projects.

### Future Directions

SimTracker can be further developed to extend its functionality and accessibility. A major step forward will be to design a Python-based version of SimTracker that can integrate seamlessly with a Python-based version of the NEURON code template. This upgrade will reduce the necessity for reading and manipulating text files; instead Python variables can be passed between SimTracker and NEURON. Further, SimTracker or parts of its functionality can then be used in conjunction with other Python-based tools such as those found at Neural Ensemble (http://neuralensemble.org/).

Future upgrades of other simulation management tools may also incorporate more functionality of SimTracker as its utility becomes apparent and modelers and experimentalists grow accustomed to increased transparency, organization, and documentation in models and model code. Additional subtemplates of the NEURON code template can be created for specific types of models. For example, some model templates that would be broadly useful and easy to adapt from the template included here are 1) a template that reads in experimental spike train data to stimulate a model network or 2) a template that incorporates more sophisticated synaptic mechanisms into its connections, such as short or long term plasticity. Existing published model code can also be reorganized into this model code template so that it can be used with SimTracker.

Not only are reviews and special issues of journals working to maintain a cohesive view of literature but individuals and institutions are also putting together databases or repositories of published data from which anyone can search or export data (http://hippocampome.org, http://neuroelectro.org,) (Ascoli et al., 2003; Tripathy et al., 2014). Downstream of the data generation and organization, modelers draw on the implications of these experiments and combine experimental observations in novel ways to aid in our goal of producing a coherent view of the anatomy and function of neural networks (Ascoli et al., 2003; Churchland and Abbott, 2016). SimTracker is well poised to serve as a resource to support closer cooperation between experimentalist and detailed network modelers.

## Methods

We developed the NEURON code template and SimTracker tool iteratively, while ensuring that the NEURON code could always be ran independently of the SimTracker to accommodate a greater range of use cases. We wrote the NEURON code template in hoc and NMODL code, and designed the SimTracker tool in MATLAB using GUIDE and custom programming in m-files (MATLAB code files). We tested the scripted and compiled versions of SimTracker on Windows, Mac, and Linux, using multiple versions of MATLAB (2014a, 2015a&b). The stand-alone versions of SimTracker were compiled on the particular operating system for which they can be used to ensure proper compatibility. We have also given users the option of downloading all relevant software at once in a complete package, by using Docker to bundle the installation of SimTracker with our model NEURON code templates, the latest version of NEURON, and the necessary terminal commands used by SimTracker.

The NEURON model code templates are available at ModelDB (Bezaire, 2016a) as well as on BitBucket at https://bitbucket.org/mbezaire/ca1 and https://bitbucket.org/mbezaire/ringdemo.

We will support SimTracker via the online NEURON forum *Tools>SimTracker* subtopic at https://www.neuron.yale.edu/phpBB/viewforum.php?f=42 and also through the issue tracker on BitBucket SimTracker repository at https://bitbucket.org/mbezaire/simtrackercode. Interested users can subscribe to updates for either the NEURON forum area or the BitBucket repository. Documentation and information of general interest for SimTracker users is available in the 15 tutorials within this work, with screenshots for Tutorials 1-5 found in Figures 8-19, as well as online in a variety of formats. Those who need further assistance with SimTracker are invited to join and post online at the NEURON forum, within the ‘tools’ section that is regularly monitored by us for SimTracker queries.

We have listed SimTracker in a variety of locations to increase its visibility, including SimToolDB (Bezaire, 2016b) and anywhere our model code is available or documented. The latest SimTracker code is always available on BitBucket. We have set up a public repository where anyone can access the code. SimTracker is covered by an MIT license so that others can use it as they see fit, while ensuring that it will remain freely available in the future, as required by the Open Source Definition (https://opensource.org/osd).

To aid with the display of model components, we created the WebsiteMaker function within SimTracker. The WebsiteMaker works in conjunction with a website template that we created. The website is implemented using HTML, CSS, Javascript and HighCharts for the graphs within the site (Figure 20). It allows users to browse the components of the model and view the experimental characterizations produced by SimTracker.

**Figure 20:**
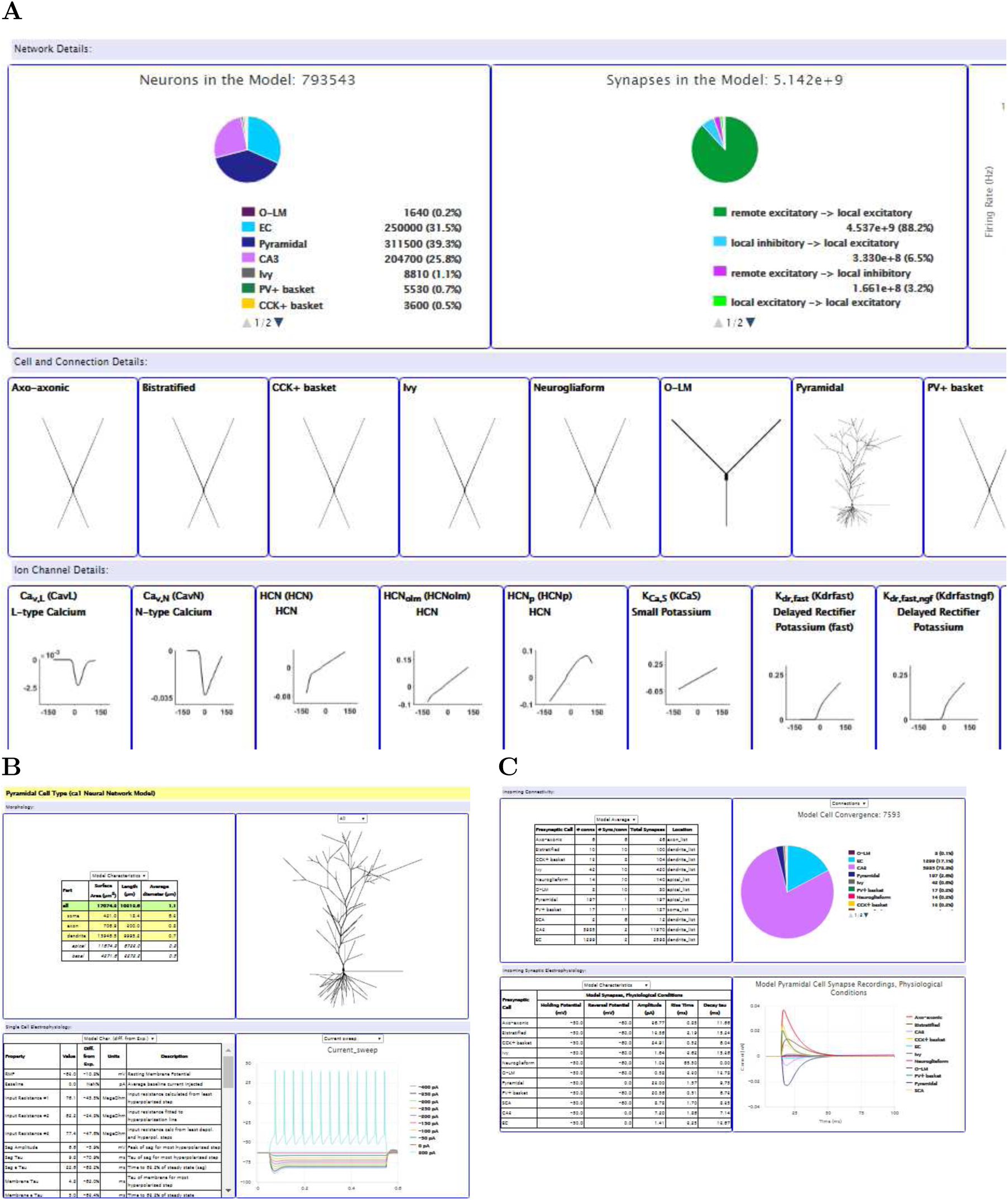
Website Template: The website template allows users to look at the characterization of model components in experimental terms as well as compare the properties with the experimental data used to constrain them, to check the constraints. (A) The home page of the website. Clicking on a cell type or channel type opens a new page with additional details such as those shown in B – C.

## Acknowledgements

This work was funded by the National Institutes of Health, grant numbers NS35915 and BRAIN Initiative NS90583 (to I.S.), F32-NS090753 (to M.B.), and R01-NS11613 (to NEURON developer Michael Hines), as well as the National Science Foundation, grant number DGE-0808392 (to M.B.) and 1458495 (to NEURON developer Ted Carnevale). This work used the Extreme Science and Engineering Discovery Environment (XSEDE), which is supported by National Science Foundation grant number ACI-1053575. Specifically, the National Science Foundation provided supercomputing resources via an XSEDE Research Allocation to I.S. (TG-IBN140007) and XSEDE Startup Allocations to I.S. (TG-IBN130022) and M.B. (TG-IBN100011) and via the Neuroscience Gateway with the support of NSF grant 1458840 (to Majumdar et al.). The authors acknowledge the Texas Advanced Computing Center (TACC) at The University of Texas at Austin for providing high performance computing resources that have contributed to the research results reported within this paper (http://www.tacc.utexas.edu). Parallel supercomputers used in this work include: Stampede and the retired Ranger, owned by the University of Texas’ Texas Advanced Computing Center (TACC); Trestles and Comet, owned by the San Diego Supercomputing Center; University of California at Irvine’s High Performance Computer and the retired Broadcom Distributed Unified Cluster.

We would like to thank NEURON developers Michael Hines and Ted Carnevale for their consistent, immediate, and precise support especially with the initial development of the parallel NEURON code but also throughout the entire project. We also thank the TACC team, the SDSC team (especially Glenn Lockwood, Amitava Majumdar, Subhashini Sivagnanam, Mahidhar Tatineni, and Kenneth Yoshimoto), and UC Irvine’s HPC team (especially Joseph Farran and Harry Mangalam) for their excellent technical support throughout this work. We would also like to thank Padraig Gleeson, Andras Ecker, Tom Morse, and Jeff Teeters for their assistance. Finally, we thank Jesse Jackson and Sylvain Williams for allowing us to include their spectrogram analysis script within SimTracker, and the Chronux team for allowing us to include some of the Chronux functionality within SimTracker.

## Tutorials

Here, we show how to use SimTracker to run NEURON simulations. The first five tutorials are meant as a basic introduction to SimTracker and include screenshots to guide the user through each process. They cover: 1) installing SimTracker, 2) setting up a new model project, 3) designing and executing a simulation, 4) analyzing simulation results, and 5) organizing simulations. For those who want a straightforward introduction to the most generally useful SimTracker components, these tutorials should be sufficient.

The next tutorial, Tutorial 6, is applicable for both modelers and experimentalists. Modelers can use it to extract cell properties from raw experimental data to serve as model cell constraints. Experimentalists can use it to automate their analysis of cell properties from AxoClamp files, increasing the efficiency and consistency with which they process their data.

The remaining tutorials are specific to particular functions within SimTracker, become more technical in scope, and do not contain pictures. For users who want or need to install SimTracker and its components onto their computer individually, rather than as part of a Docker package, tutorial 7 walks through the steps necessary for a complete, manual install. Tutorials 8-10 demonstrate how to use SimTracker to run large scale parallel NEURON models on supercomputers. The remaining four tutorials highlight the functionality of other tools within SimTracker: 11) how to manage parameter sets within SimTracker, 12) how to characterize model components using the CellClamp tool, 13) how to perform network clamps (Bezaire et al., 2016) using the Network Clamp tool, 14) how to use NEURON’s Multiple Run Fitter to tune network components, and 15) how to display the model description and results on a website.

Finally, all SimTracker functionality is open to customization. For users who want to customize SimTracker or its outputs, review the MATLAB code, or even run the NEURON code independently of SimTracker, we recommend consulting the instruction manuals for SimTracker or the NEURON template at http://mariannebezaire.com/simtracker/. Additional training is also available online.

While working through the tutorials, keep in mind that steps that include the text `terminal-$ enter_this_command` mean that, at the user’s local terminal, the text after the dollar sign should be entered (in this example, `enter_this_command`). Any step that advises entering “text” somewhere means that only the characters within the quotes, not the quotes themselves, should be entered.

### Tutorial 1: Install SimTracker

There are three ways to install SimTracker: 1) using Docker, a virtual software container which has a preconfigured software environment that includes all software necessary to run SimTracker and NEURON in Linux, 2) by installing SimTracker on a system that has MATLAB, NEURON, and Mercurial, and 3) by installing a stand-alone version of SimTracker that doesn’t require a separate MATLAB license on a system with NEURON and Mercurial. In this tutorial, the user will use the first approach on a Linux computer to create a fresh instance of a Docker container with SimTracker installed.

1. To create a fresh container instance with SimTracker installed, install Docker either by downloading it from https://www.docker.com or by using the package manager corresponding to the Linux distribution on the computer. Docker installation instruction for different Linux distributions are available at https://docs.docker.com/engine/installation/linux/.
2. After installing Docker, download and install the SimTracker ringdemo image with the following command:

~~~
terminal-$ docker pull solteszlab/simtracker
~~~ This will download the SimTracker Docker image and save it on the computer.
3. Next, create a ‘repos’ directory which will be used for storing model code and results data generated when running simulations:

~~~
terminal-$ mkdir repos
~~~
4. Then, start the SimTracker container by entering the following command:

~~~
terminal-$ docker run –rm -it -e DISPLAY=$DISPLAY -v /tmp/.X11-unix:/tmp/.X11-unix -v $PWD/repos:/home/docker/repos solteszlab/simtracker
~~~ Note: The options `-e` and `-v` ensure that GUI applications based on the X11 standard can run in the Docker container and their graphics can appear on the host computer. The second `-v` option mounts the newly created ‘repos’ directory as a volume inside the Docker container. It is important to mount this directory from the host computer, making it available within Docker. Unless special steps are taken, files and data saved to other directories within the Docker package, which are not accessible outside of Docker, will also not be available even within Docker after closing and reopening the Docker package.
5. Finally, run SimTracker with the following command:

~~~
terminal-$ ./simtracker/run_SimTracker.sh
~~~ SimTracker will start and ask for a repository directory. The next tutorial explains the steps necessary to create a new repository. When done using SimTracker, the Docker container can be exited with the ‘exit’ command. Please consult the online Docker documentation at https://docs.docker.com/engine/userguide/containers/dockerimages/ for further details on how to use Docker images and software containers, including use of the ‘docker commit’ command to save data and settings within the package.

### Tutorial 2: Create a new model repository using SimTracker

Now, Tutorial 2 will show how to set up a new model (Figure 8), which will be an occasional task. To create or add a model repository to SimTracker, do the following:

1. From the File menu, choose ‘New Repository’ (Figure 8A).
2. Because the repository directory is empty, SimTracker will offer a choice of downloading a sample repository from BitBucket, such as the ringdemo network model code at https://bitbucket.org/mbezaire/ringdemo. Click the ‘ringdemo’ button to download the code as the remaining tutorials in this section will use the ringdemo network (Figure 8B). Note: if users choose not to download code because, for example, they are adding a model repository that already contains its own code, they will instead be given the option for SimTracker to perform a check of the files in the repository and add any missing ones to complete the NEURON template used with SimTracker.
3. SimTracker will download the code and register the repository, then display a message when finished (Figure 8C) and will now list the path to the repository at the top of the SimTracker window (Figure 8D). Note: If users choose not to download the sample code, at this point a text file would open containing a to-do list of any missing files and folders the repository requires. Make sure to address each item on the to-do list before continuing to work with the new model repository. The model repository is now ready. The next tutorial will walk through the processes of creating, executing, and analyzing a simulation, tasks that users will perform many times.

**Figure 8:**
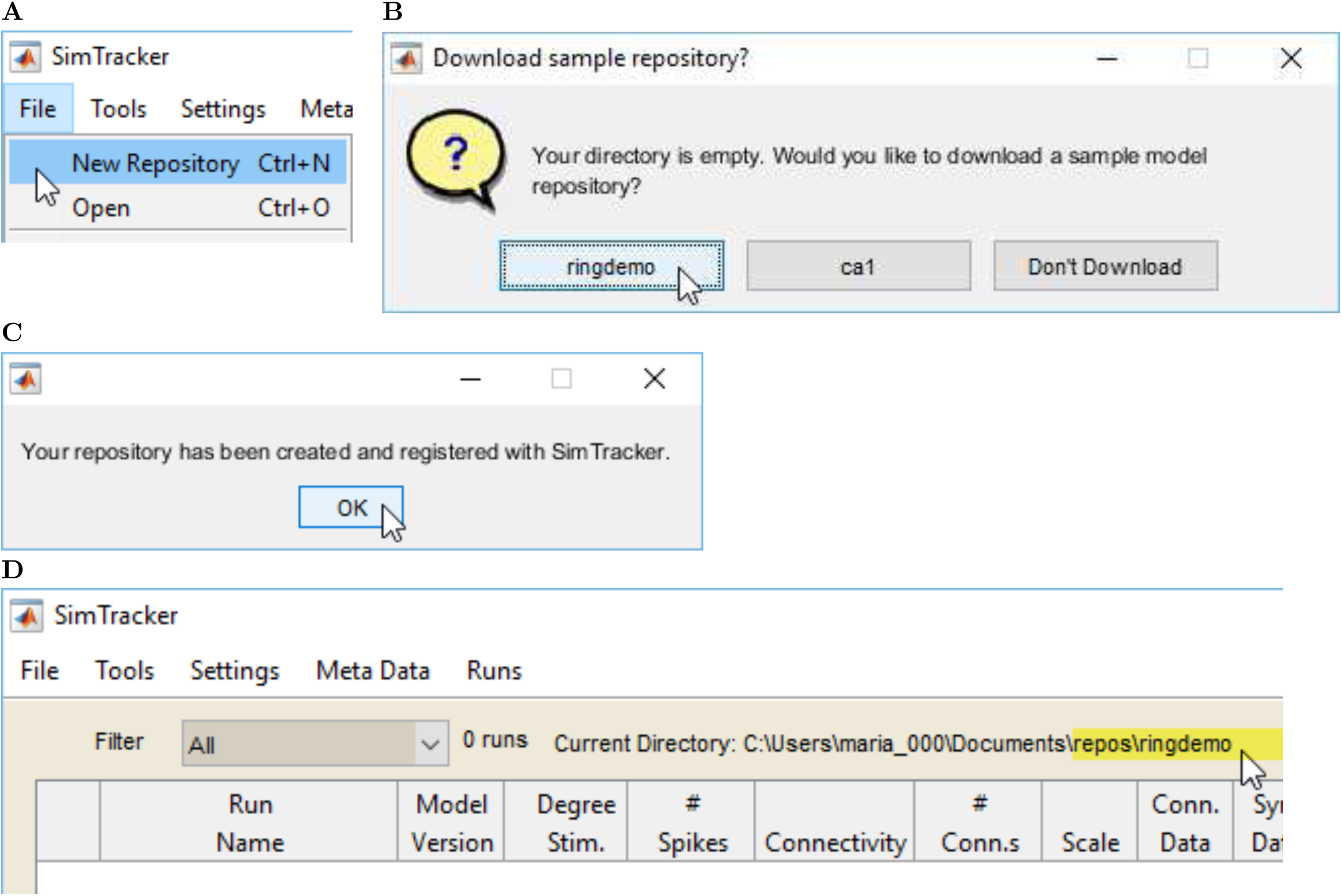
Tutorial 2: How to set up a new model code repository. The steps of this tutorial will need to be completed each time the user creates a new model for use with SimTracker. 8A New repository command within File menu, 8B prompt to download sample model code, 8C message confirming repository creation, 8D path to repository displayed on SimTracker window.

### Tutorial 3: Design, execute and upload a simulation

This tutorial covers designing and executing simulations, which will be a frequent task. Before designing specific simulations for a model, users will likely want to create some datasets to specify the cells and size of the model, the numbers and weights of its connections, and the kinetics of its synapses. However, those steps are not necessary for this example ringdemo network because those data sets have already been created and are included in the download; therefore the steps are detailed in a later tutorial. This tutorial will simply show how to view the datasets that have already been created, for the purpose of understanding all the components of the model before designing and executing a simulation (Figures 9-11).

1. Within the ‘Tools’ menu, choose the ‘Cell Numbers’ option (Figure 9A).
2. A dialog box will open, allowing users to view the Cell Numbers datasets associated with this model repository.
3. From the popup menu at top, choose number #100 to display the dataset that will be used to set the types and numbers of cells in the ringdemo network (Figure 9B). The table will refresh, showing two rows with information about the two cell types in our model (Figure 9C). The left-most column lists the name users have given the cell in their model. The next column lists the name of the cell template file to use for this cell, from the list of cell template code files in our model repository. The third column gives the number of that cell type to include in the model. The last column is only relevant for 3D models with layers, listing the layer in which that cell type is found (starting from layer 0). After viewing, close the window.
4. Next, from the ‘Tools’ menu, choose ‘Connections’ (Figure 9D). From the popup menu at top, choose number #100 to display the connectivity dataset to use (Figure 9E). A matrix will appear, including all the cell types in the network. Down the left side of the matrix (the row labels) are given the presynaptic cells, while the top row (the column labels) gives the postsynaptic cells. At the intersection of each presynaptic and postsynaptic cell, there are entries for the average number of connections from all cells of that presynaptic type onto one postsynaptic cell of that type (aka, the convergence), the number of synapses comprising one connection, and the weight of each synapse. After viewing the connection information, close the connection tool.
5. Finally, from the ‘Tools’ menu choose ‘Synapses’ (Figure 9F) and a table will appear. From the popup menu at the top left, choose dataset #100 to view the synapse data for the model (Figure 9G). Only one postsynaptic cell at a time will be displayed; choose from the pop-up menu at the top middle of the table (Figure 9H) to specify which post-synaptic cell to view. Then the table will display the possible presynaptic cell types for that postsynaptic cell, along with entries for the rise and decay time constants of the synapse and the reversal potential of the synapse (Figure 9I). When finished, close the window. The next steps show how to design a simulation that uses these datasets.
6. To design a new simulation, from the Runs menu choose ‘Design Run’ (Figure 10A). The property fields in the SimTracker will become editable (Figure 10B).
7. Enter a unique name for the simulation (allowed characters include letters, numbers, and the underscore character), such as MyFirstRingDemo (Figure 10B).
8. Optionally, enter comments about the simulation. It is often helpful to users if they note the specific question or idea that triggered them to run this particular simulation.
9. Specify the code version to use for this simulation. Usually, this will be the most recent version of the code. The list of available code versions will correspond to each time users ‘committed’ a version of the code in their model code repository and will also list the comments they entered when committing that code version. Select the most recent version, #9 (Figure 10C).
10. Specify the machine to use for running the simulation. For users to run this simulation on their own computers, they must find the unique name of their computer from the list and select it (Figure 10D). Then, in the field to the right of the machine name, specify the number of processors to be used to execute the run. On a personal computer, users can either set this to 1 or, to run the network in parallel, set it to the total number of cores (aka processors) on their machines.
11. Select from the various menus for simulation control to specify the stimulation, connectivity algorithm, cell numbers, connection numbers, and synapse kinetics for the model. For this ringdemo network, choose the following settings (Figure 10E):
  - Stimulation: pulse
  - Connectivity: ring
  - Cell set: 100
  - Connection set: 100
  - Synapse set: 100
12. In the table on the right side of the SimTracker, specify additional properties for simulation control, including the scale (size) of the model network, the length of time the simulation should last, the temporal resolution, whether to print specific results files, and many other options (the options that appear in this table can be customized by choosing ‘Parameters’ from the ‘Settings’ menu). Because this ringdemo network is so small, it can run it at full scale. Set the following parameters and leave the remainder set to their defaults (Figure 10F):
  - Scale: 1
  - Sim. Duration: 101
  - Temporal Resolution: 0.025
13. Upon finishing the specification of simulation properties, click the ‘Save Run’ button that appeared when upon starting to design the simulation (Figure 10G). The simulation will now appear in the list of simulations at the top of the SimTracker. The property fields will become uneditable (to change the run prior to execution, choose ‘Edit Run’ from the ‘Runs’ menu). The process for executing a simulation differs somewhat depending on whether users are executing it on their personal computers, using the NSG portal, or accessing a supercomputer independently of the NSG portal. The following list details the process for using a personal computer. For instructions on using the NSG portal or a supercomputer directly, see Tutorial 10.
14. To execute the simulation, select its row from the table at the top of the SimTracker, and then from the ‘Runs’ menu choose ‘Execute Run’ (Figure 10H).
15. The computer’s CPU may become very active and users who are using MATLAB scripts to run the SimTracker, will see the MATLAB status bar display ‘Busy’ until the simulation completes (Figure 11A). Additionally, a NEURON window may pop up (Figure 11B).
16. When the simulation finishes, a dialog box will appear that states ‘The run has been executed’ (Figure 11C). Once this message appears, continue to the next step.
17. Select the simulation in the SimTracker (make sure to view the ‘Not Ran’ view) and, from the ‘Runs’ menu, choose ‘Upload Run’ to display the results of the run in the SimTracker (Figure 11D).
18. Once the simulation is successfully uploaded to the SimTracker, a dialog box will appear stating that ‘Results have been uploaded.’ (Figure 11E) The SimTracker table will switch to the ‘Ran’ view (Figure 11F). In both the list of runs and the form view below, SimTracker will show the executed simulation’s run time, number of spikes, and execution date (Figure 11G). After the simulation has been executed and uploaded, it is time to analyze the results (Figure 12).

**Figure 9:**
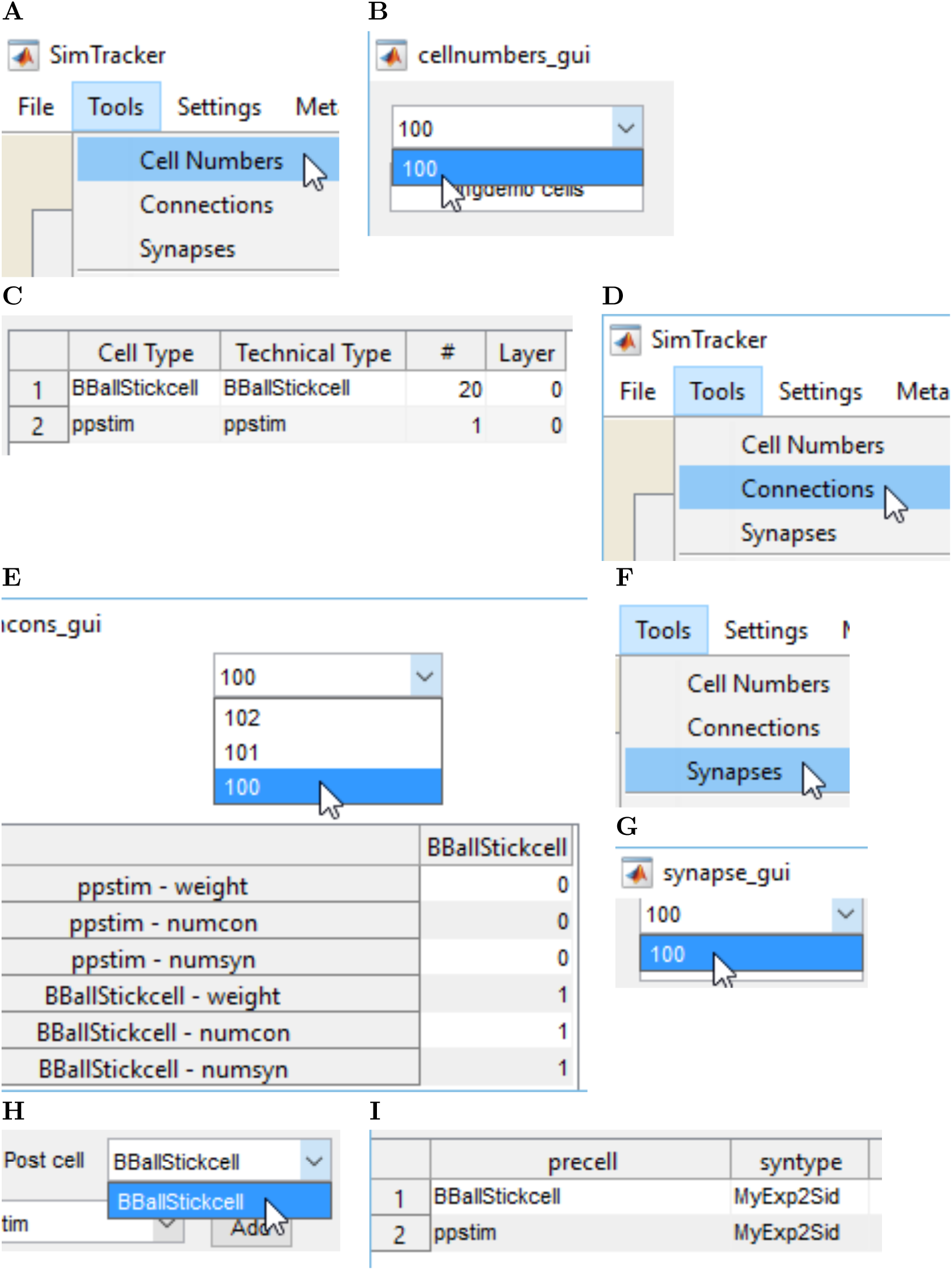
Tutorial 3: How to view, edit, and make new datasets for use in simulations. (A-C) Viewing the Cell Numbers datasets. (D-E) Viewing the Connections datasets. (F-I) Viewing the Synapses datasets.

**Figure 10:**
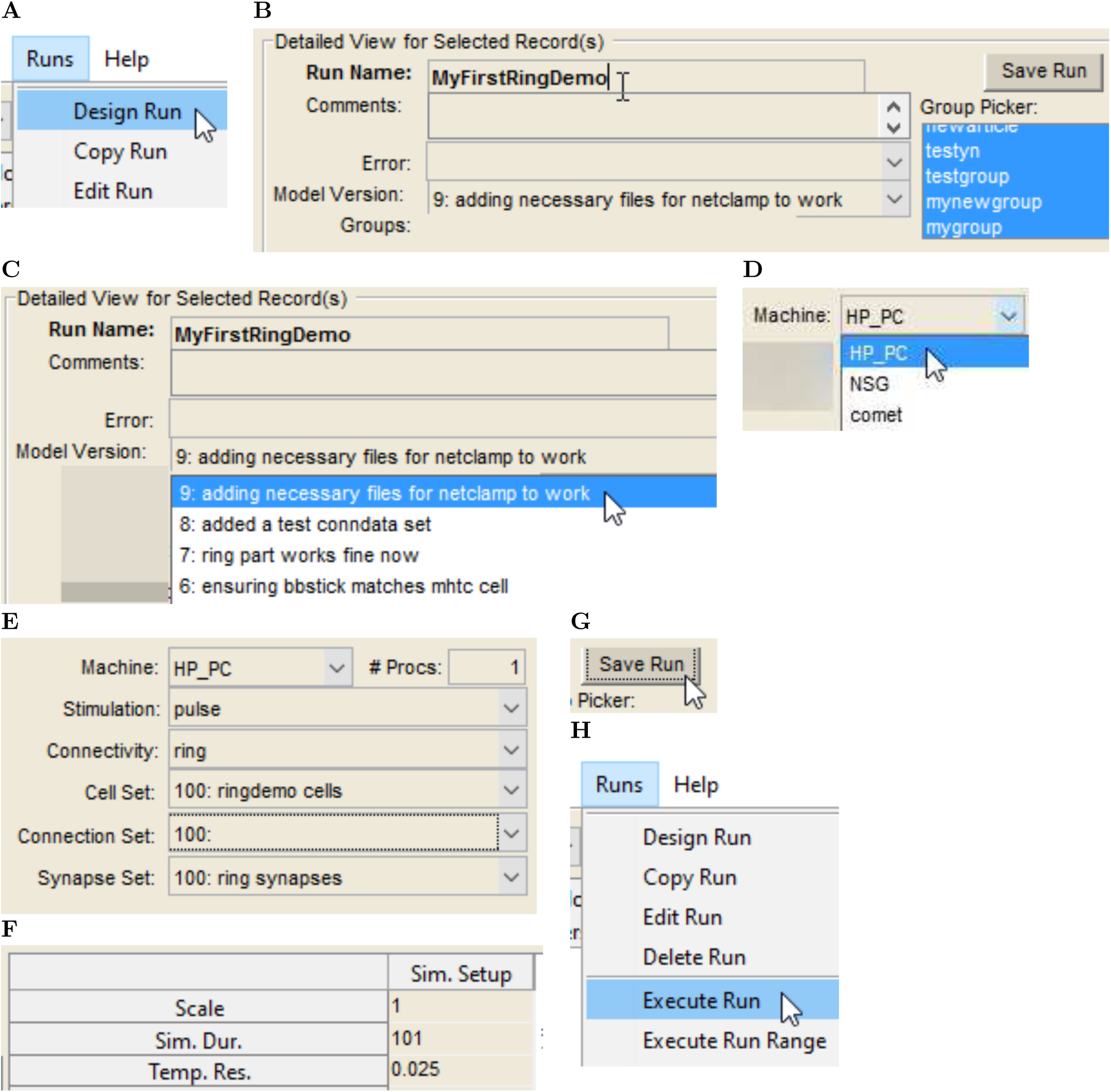
Tutorial 3, continued: Designing and executing a simulation run. (A-F) Create a new simulation and setting its parameters, including a unique Run Name. (G) Save the simulation after setting desired parameters. (H) Execute the simulation run after saving it.

**Figure 11:**
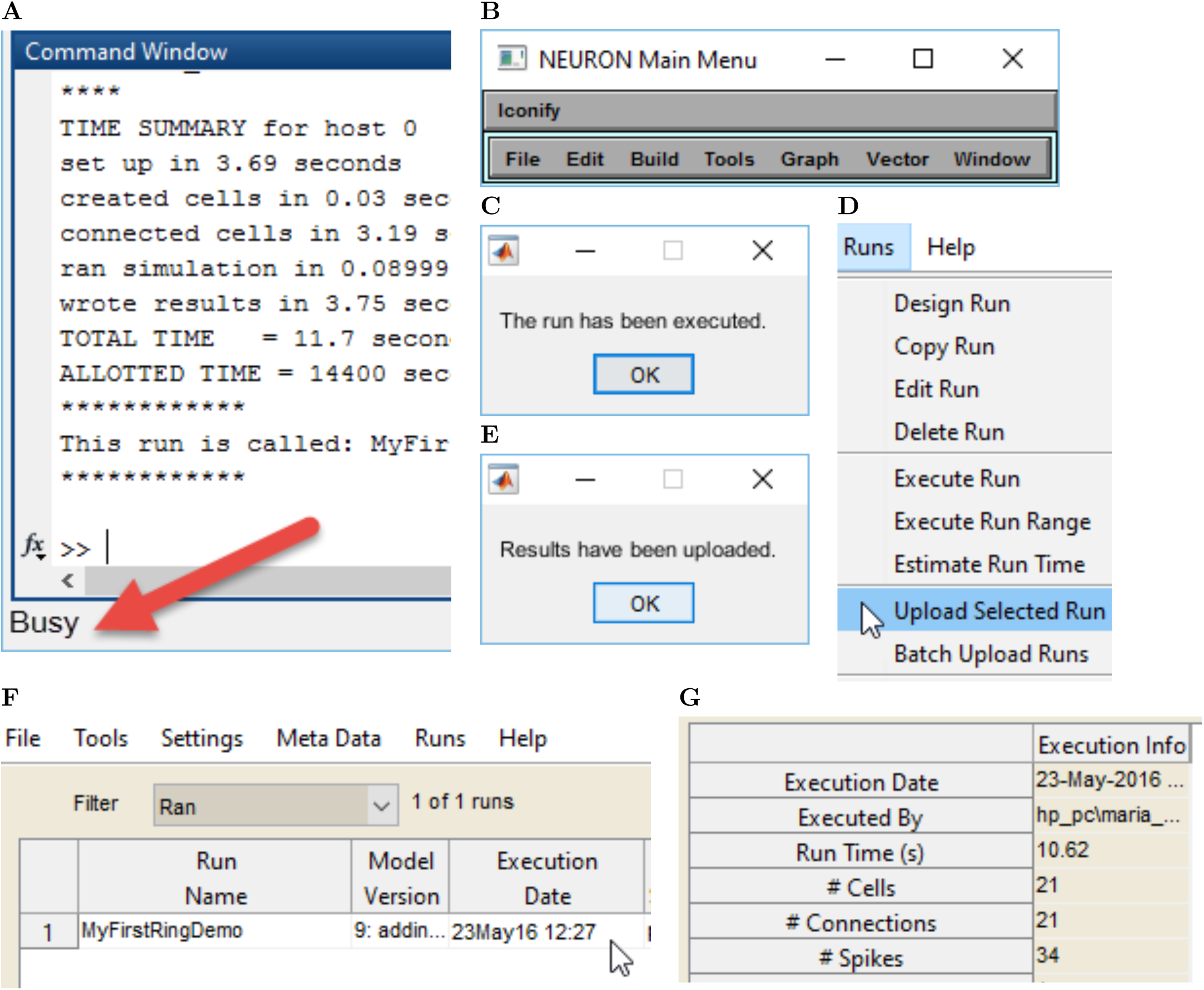
Tutorial 3, continued #2: Execution of simulation and uploading of results. (A-B) Indications that NEURON is running a simulation on the user’s computer. (C) After a locally-run simulation has completed, this dialog box will appear. (D) Upload to SimTracker results from a completed simulation. (E-G) When the results have uploaded to SimTracker, a dialog box will appear and the execution info table within SimTracker will populate with additional properties of the simulation.

**Figure 12:**
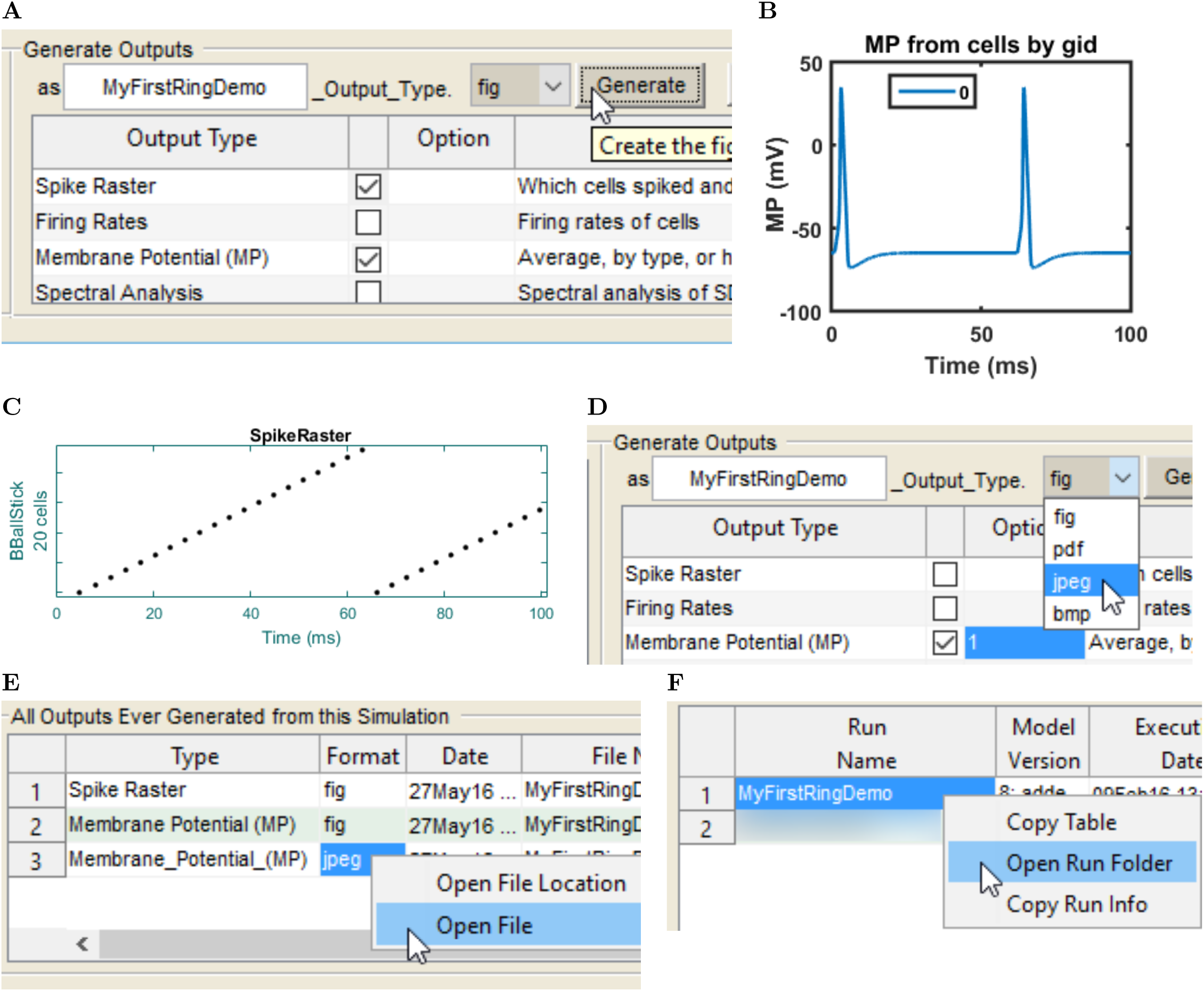
Tutorial 4: Analyze a completed simulation. After simulation results have been uploaded, a variety of analyses are available to characterize the model and its behavior. (A) Select the desired output from the table and click the Generate button to produce the output. (B-C) Possible simulation outputs include (B) a membrane potential trace and (C) a spikeraster.(D) Save image files of analysess in the results directory of the simulation. (E-F) There are two methods of accessing the saved image file in the simulation’s results directory: right-clicking (ctrl+click on Mac) either (E) the Outputs Generated list or (F) the Simulation Run list.

### Tutorial 4: Analyzing a simulation

Any simulation that has been executed and uploaded to the SimTracker is available for analysis. This tutorial walks through how to use some standard analyses available within SimTracker. However, users may also access the results from each simulation directly, by opening the directory with the same name as the simulation, found within the ‘results’ directory of the model repository. The results directory can also be opened by right-clicking (Ctrl-click on Mac) the entry for the simulation in the main SimTracker runs list. The spikeraster.dat file within the results folder contains a list of all spike times and which cell caused each spike during the simulation. The ringdemo simulation produces a spikeraster file that contains the times found in Table 1, which compare well to the results from Hines and Carnevale (2008), found in their section *3.3.5 Reporting simulation results*.

**Table 1:**
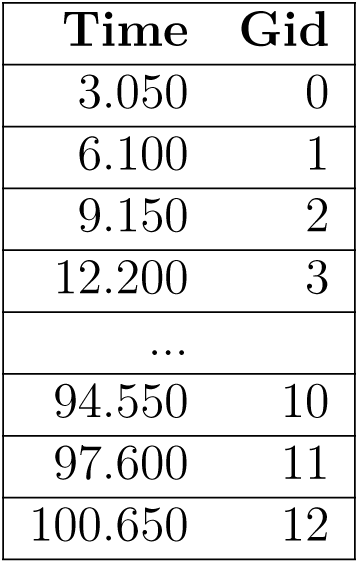
Beginning and end of sorted spike raster file for our implementation of the Hines and Carnevale (2008) ring demo network. Due to a different conduction delay parameter in our network, our ring cell spike times are delayed 1 ms relative to the results of Hines and Carnevale (2008); however, the order and relative delays of the spike times are all equal to the output of the original ringdemo network.

1. Select the completed simulation of interest from the table in the SimTracker.
2. Look at the table on the bottom right side of the SimTracker for a list of available outputs for that particular simulation (the list will vary depending on which output files users choose to print for their simulations). For now, click to select the checkboxes next to the ‘Spike Raster’ and ‘Membrane Potential’ (Figure 12A) outputs. Note: For more information on any particular output, click on that row to highlight that output type and then hover the mouse anywhere over the output table to see a tooltip with guidance about the selected output type.
3. After selecting the outputs of interest, click the ‘Generate’ button to produce them as MATLAB figures. If prompted to enter a GID (global ID number for a cell type), enter ‘0’ to display the membrane potential for cell #0. Users can zoom in, pan, and otherwise edit or save the figures once the MATLAB figures are generated (Figure 12B and 12C).
4. Alternatively, users can directly export the figures to an image format of their choice by selecting the ‘fig’ menu option in the file extension pop-up menu and choosing a different file format (such as jpeg, bmp, pdf, etc.) (Figure 12D).
5. Exported images will be saved in the results folder for that particular simulation, which will be logged in the figure-generation log on the lower left side of the SimTracker (Figure 12E). To view the file location of the saved image, users can right click in the figure-generation log or in the main simulation list window and choose to open the simulation results directory (Figures 12E and 12F).

### Tutorial 5: Organize simulations

After completing many runs, users may wish to organize them in various ways. One possibility is to add labels to the runs as well as adding searchable comments. Users can set the view filter to only show runs with (or without) particular labels, as well as runs with similar names, or runs meeting a custom search requirement of their choice. Users can and should back up their runs (both configurations and results sets) using SimTracker. Archive runs that are no longer needed (deleting executed runs from the SimTracker is not recommended). Users can also, export and import runs to share with others. This tutorial will illustrate labeling, filtering, and archiving a completed simulation, then importing it back into SimTracker. To prepare for this tutorial, please try running an additional simulation following the steps in Tutorial 3, giving it a unique name. After uploading the results, proceed with this tutorial (Figures 13 – 14).

1. First, add a grouping category to SimTracker by going to the Settings menu and choosing Groups (Figure 13A).
2. When the group list appears, click the ‘Add line’ button and in the left cell of the new line that appears, enter a group name ‘testgroup’ (Figure 13B). Then click the ‘Save’ button and close the window. The entry ‘testgroup’ will now appear in the list of groups on the SimTracker.
3. Next, ensure that the view is set to ‘Successfully ran’ so that the completed runs show. Then, click the MyFirstRingDemo entry in the table to select it.
4. Next, find the ‘testgroup’ entry in the group list and click it. The label ‘testgroup’ should appear in the ‘labels’ section of the simulation entry (Figure 13C, highlighted section).
5. Now, from the view filter menu, choose ‘Group A,B,C filter’ (Figure 13D).
6. In the dialog box that appears, find the line for ‘testgroup’ and from the menu next to it, pick the equals sign entry (=), then click OK (Figure 13E).
7. Now, the only run showing in the SimTracker should be MyFirstRingDemo (Figure 13F).
8. Again select the MyFirstRingDemo run and now from the File menu, choose ‘archive’ (Figure 14A). Users can archive many runs simultaneously by selecting all desired runs before choosing the ‘archive’ command, but this tutorial will only archive one.
9. An input dialog will appear, allowing the user to enter a name for the archive. Enter a name such as ‘MyTestArchive’, and click ‘Save’ (Figure 14B). SimTracker will create an archive in the backup folder under the name specified by the user. The backup folder is automatically created in the same directory where the other repositories are stored.
10. After MATLAB has created the archive, it will ask if the user is ready to delete the archived files from SimTracker. Click ‘Yes’ and the archival process is complete (Figure 14C). Note: users who want to create a backup file of a run without removing it from SimTracker, can do so by choosing File>Backup rather than the Archive command.
11. Now, the run should have disappeared from the SimTracker. Change the view setting to ‘All’ and verify that the run no longer appears (Figure 14D).
12. Next, import the run back into SimTracker by going to the File menu and choosing ‘Open’ (Figure 14E).
13. In the dialog box that appears, click the second option and then select the saved archive from the list (Figure 14F).
14. Click OK. The run should now appear in the SimTracker table again.

**Figure 13:**
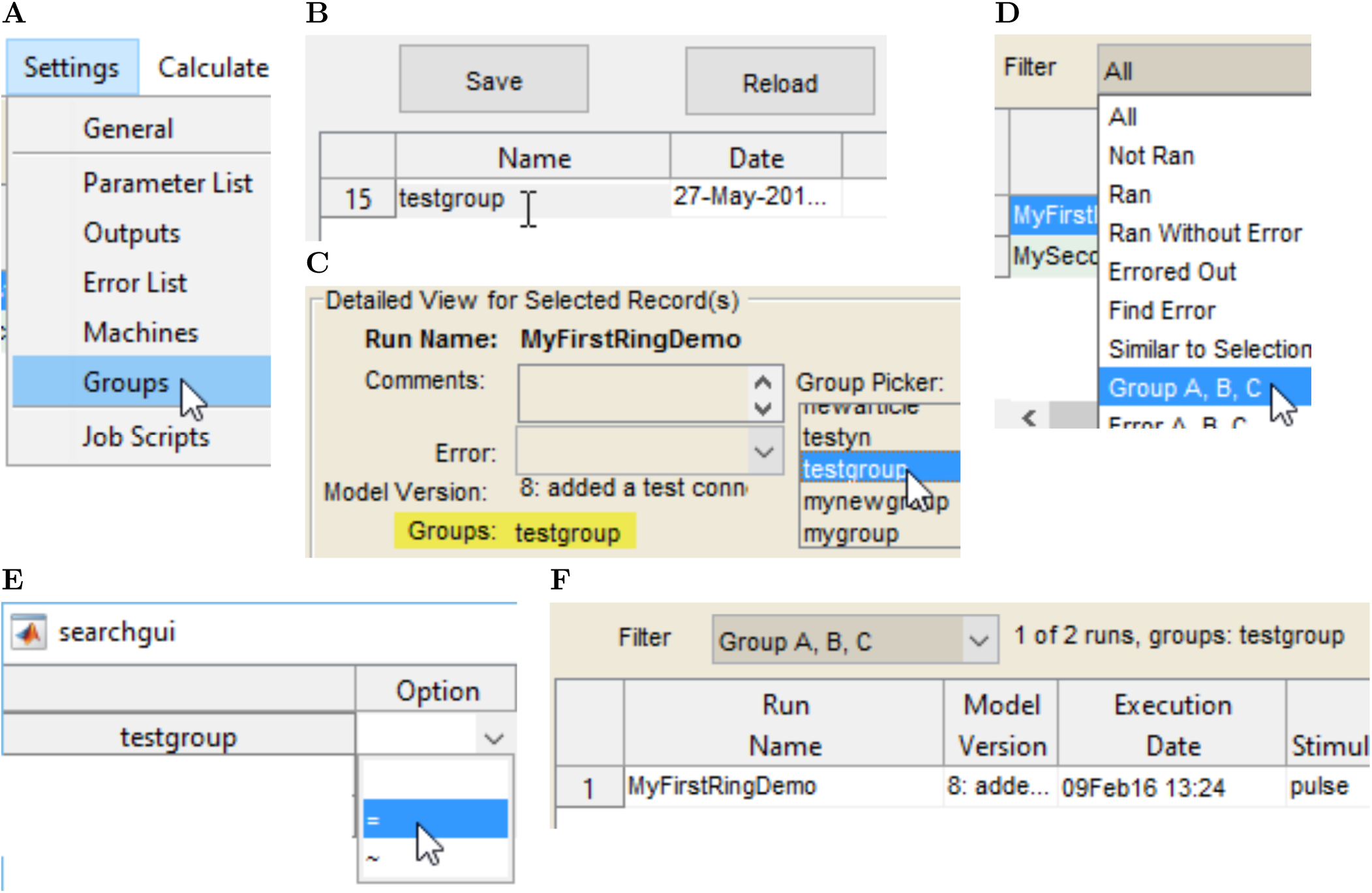
Passerine SINEs share a common ancestor and are mobilized by CR1-X.xs. Tutorial 5: How to organize simulations into groups. (A) select Groups from the settings menu.(B) enter a group discription in the box, (C) select the category to label a simulation. (D) Set the filter to show members of that group. (E) set the search criteria, (F) the view showing all simulations within the filtered group.

**Figure 14:**
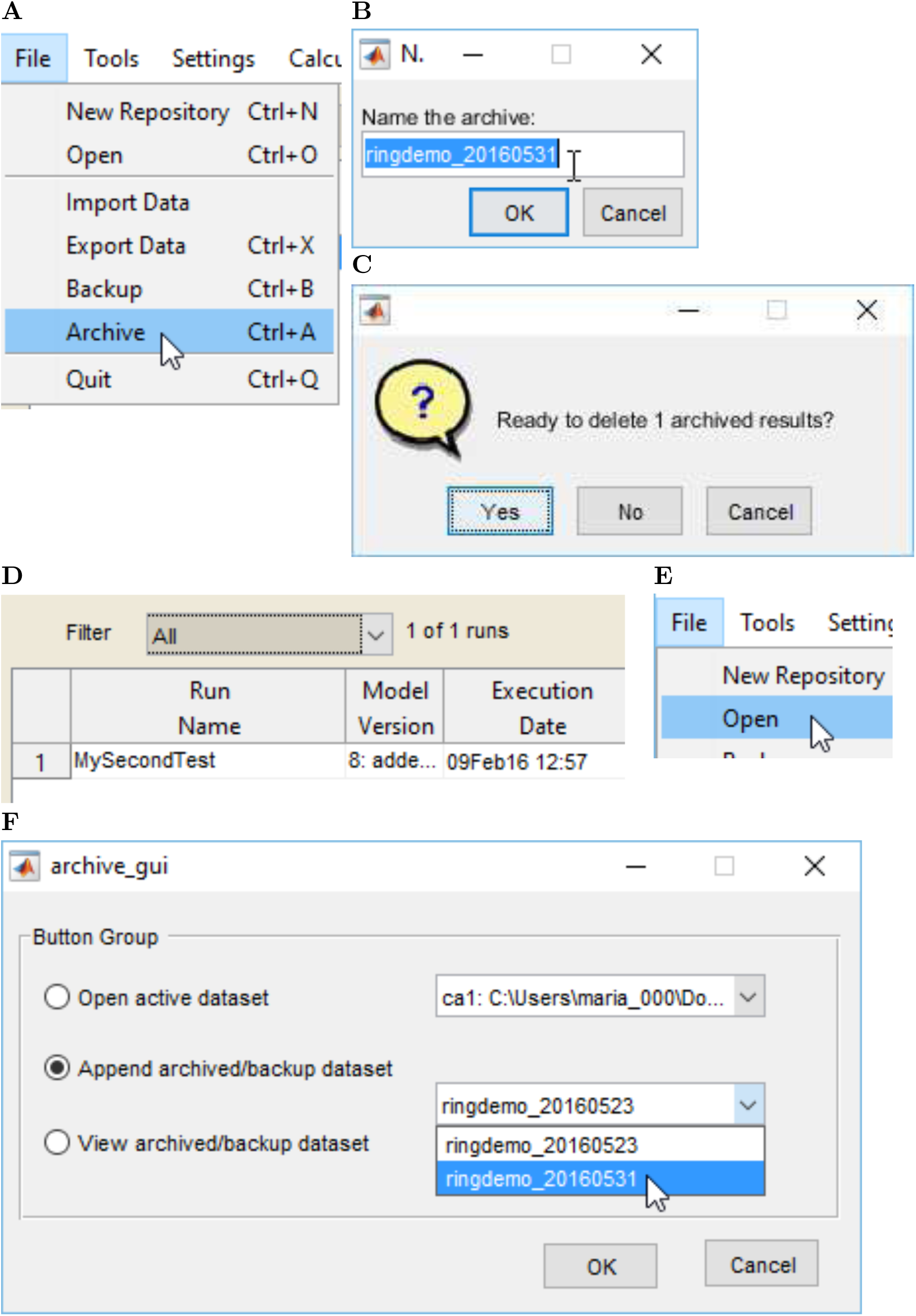
Tutorial 5, continued: How to archive and re-add simulations to SimTracker. (A) Select Archive from the File menu, (B) name the archive package, (C) click ‘Yes’ to clear the files from SimTracker, (D) observe that the archived files have been removed from view, (E) go to File and then Open (F) click the Append radio button and then select an archive to reload.

### Tutorial 6: Automatically analyze experimental data using CellData

Both experimentalists and modelers may wish to automate analysis of AxoClamp recordings from biological cell current sweeps. The CellData tool reads in AxoClamp files in binary (ABF) or text (ATF) format. If the file contains recordings from a current injection sweep, such as the protocols commonly used to characterize firing curves and intrinsic electrophysiological properties of single cells, the CellData tool will process the file to determine spike threshold points, action potential peaks, sag and steady state peaks. Then, it will allow the user to review and edit the points for accuracy (or choose a different strategy for determining the action potential threshold), before computing over 30 common electrophysiological properties of the cell.

CellData can be used independently of SimTracker. Both a compiled version and the MATLAB source code of CellData is available for download at http://mariannebezaire.com/simtracker/.

1. From the Tools menu of SimTracker, choose the ‘Experimental Comparison (CellData)’ option to open the CellData tool (Figure 15A)
2. In the CellData tool, load data from an AxoClamp file by clicking the button ‘Load AxoClamp file’ (Figure 15B) and selecting the desired AxoClamp file (in ABF or ATF format) from the file picker dialog box that appears (Figure 15C).
3. As the recording data is imported, a dialog box will appear asking for additional metadata (Figure 15D). Enter the cell type and location for the cell, using whichever nomenclature is appropriate to the user’s field of study. Also enter the name of the person who recorded the cell, for future reference. Then click OK.
4. Once the recording data is read in from the cell, CellData will display some of the traces from the file in a graph at the lower right corner of the tool. The traces displayed will include recorded membrane potential from the most hyperpolarized trace, the most depolarized trace, and the trace from the current injection at the zero level, if performed (Figure 15E). Overlaid on these traces will be the corresponding current injection plots.
5. Users should look these over and ensure that the current injection timing looks correct and corresponds to the displayed potential traces, and then click the ‘Verify Current Sweep’ button (Figure 15F). A dialog box will then appear, confirming ‘Current Sweep Verified’ (Figure 15G).
6. The lower right corner of the CellData tool will now display the some membrane potential traces from the current sweep, including the potential recordings corresponding to all hyperpolarized current injections and the most depolarized current injection (Figure 16A). However, the traces must be further analyzed to calculate the electrophysiological properties of the cell. Click the ‘Calculate Threshold’ button to start the analysis process (Figure 16B).
7. The threshold calculation process can be time intensive, so the CellData tool will display a status bar to alert the user to the progress of the calculation (Figure 16C).
8. A second window will appear, displaying one at a time the membrane potential recorded at each current injection. In this interface, users can choose from one of three possible threshold calculation strategies, visually observing how the point chosen depends on the threshold calculation used. Select ‘1’ from the dropdown menu picker to set the calculation strategy for this sample cell (Figure 16D).
9. A line will be drawn through each threshold point calculated in the displayed membrane potential trace (Figure 17A, see red arrow). Users can browse through the membrane potential recordings from each current injection level by selecting the left or right arrow buttons (Figure 17A, see orange outline), while keeping track of which level they are currently viewing by looking at the description in the upper left corner of the window (Figure 17A, see red circle). Whichever threshold calculation strategy is chosen will be used to find the threshold points in the recordings from all current injection levels.
10. When satisfied with the threshold calculation strategy chosen, push the ‘Set Threshold’ button (Figure 17B) and wait for CellData to calculate all spike threshold points from all current injection levels in the file. When the calculation has finished, the second window will close.
11. Back in the CellData window, click the button to ‘Verify … Analyze’ (Figure 17C). Another status bar will appear in CellData to signal the progress of the review step, which can take awhile for large files.
12. After the tool has identified all points of interest in the membrane potential recordings from each current step, the second window will again appear, allowing users to review and correct any point (Figure 17D). The points found by the program include (for hyperpolarized traces) the trough point of the sag potential, (for hyperpolarized and subthreshold depolarized traces) the steady state potential, (for subthreshold traces only) the transient peak depolarization potential, (for suprathreshold traces) the action potential threshold, action potential peak, and afterhyperpolarization potential.
13. Figure 17E shows various buttons available for modifying the points automatically chosen by the CellData tool. To move or delete an incorrect point, either click the corresponding action button (Figure 17E) and then select the point by clicking it with the mouse, or scroll through the table of points to the right side and select the checkbox next to the point, then click the appropriate action button.
14. As when selecting the threshold, users may scroll through the membrane potential recordings associated with each current injection level, and may correct points at each level. When finished, click the ‘Done with cell’ button to close the second window and return to the main CellData interface.
15. Now CellData will display additional buttons for displaying and exporting the analyses of the cell, as well as a table of calculated electrophysiological properties (Figure 18A) and, in the lower right corner, a graph that can be set to display several different analyses by making a selection from the drop-down menu above the graph (Figure 18B).
16. Click the ‘Make Figures’ button (Figure 18C) to display an interface where users can pick which graph types to display and the format in which to display them (Figure 18D). Figures such as those pictured in Figure 19 will be displayed.

**Figure 16:**
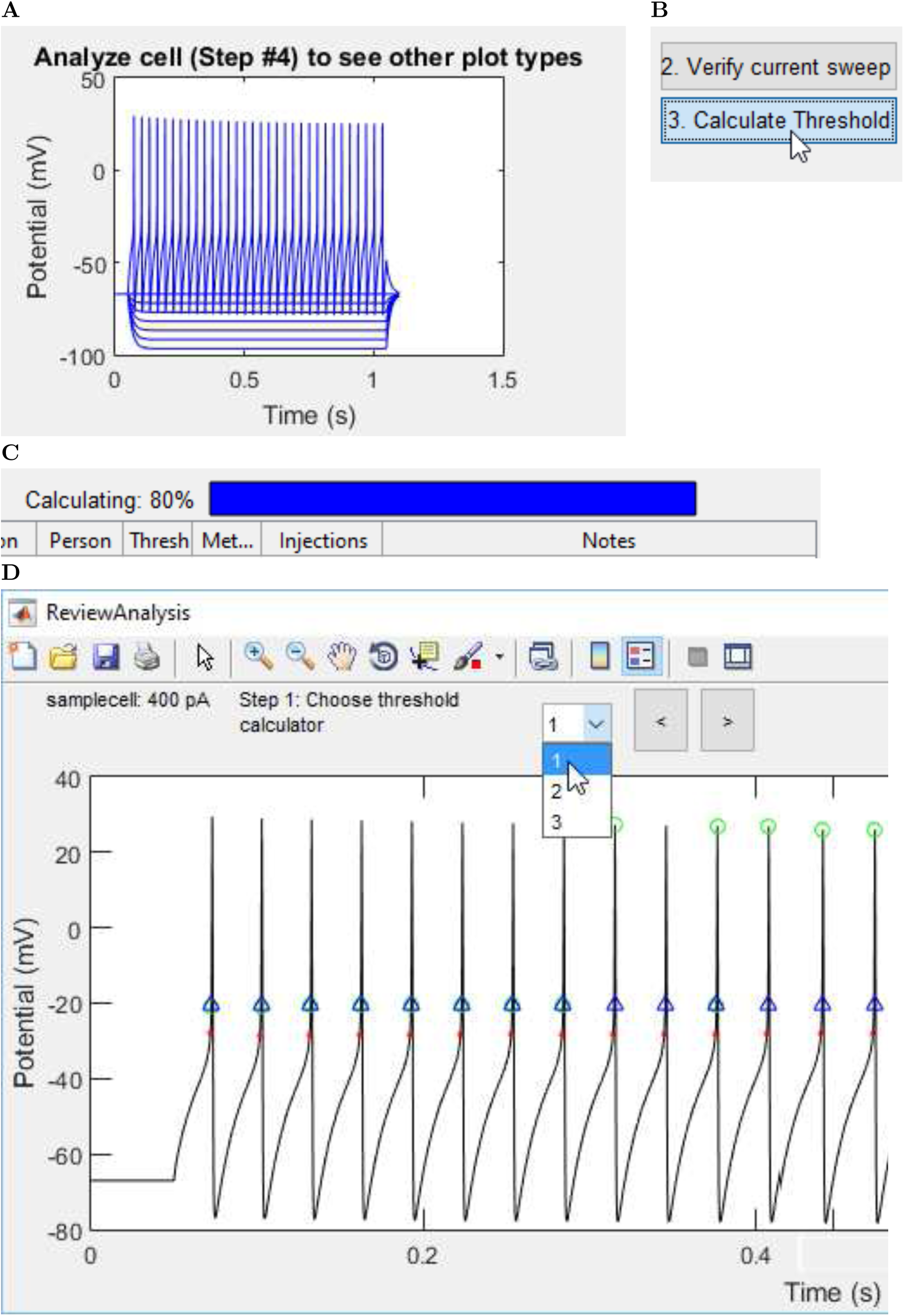
Tutorial 6, continued: How to review and export the analysis of an AxoClamp file.

**Figure 17:**
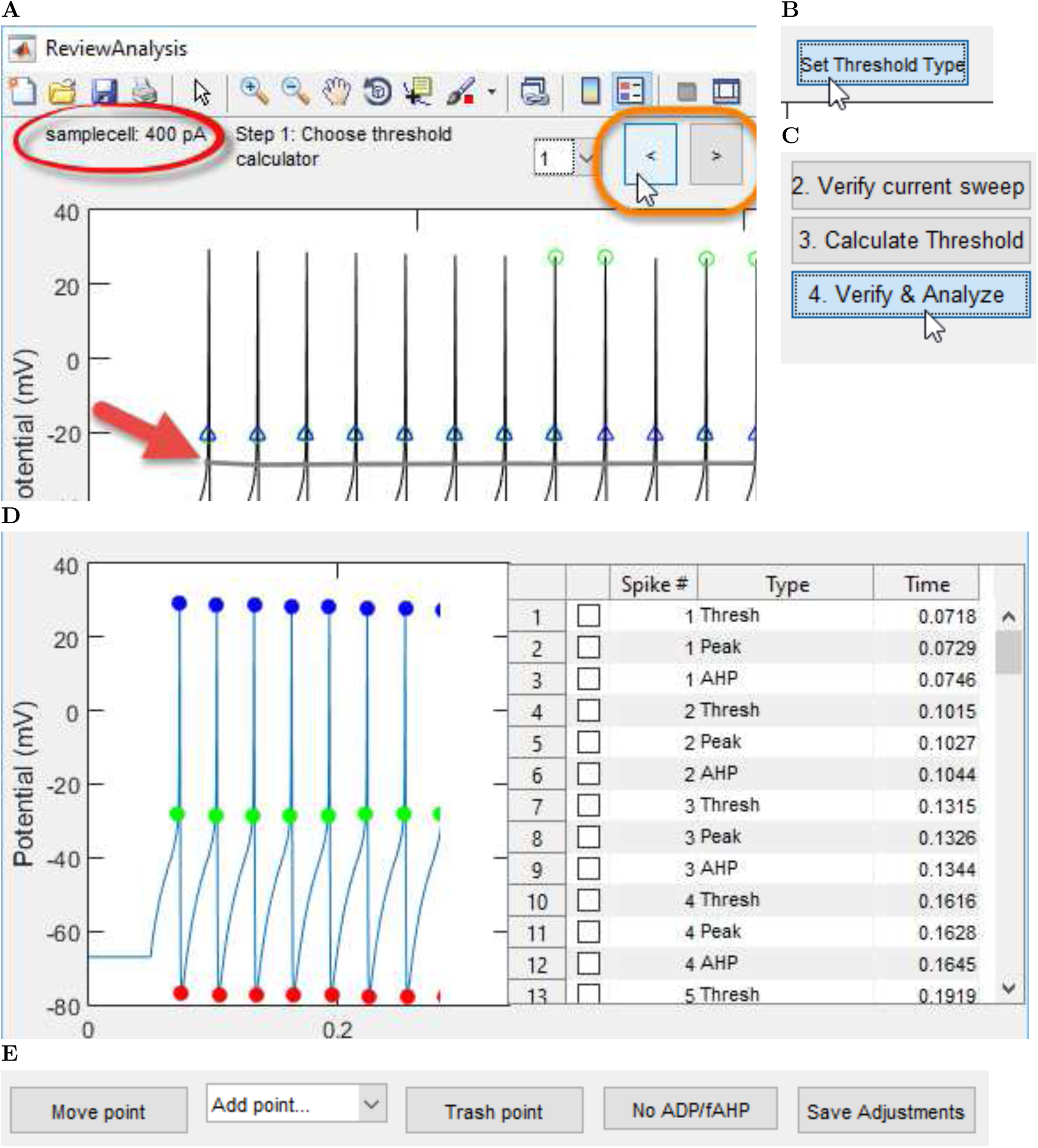
Tutorial 6, continued part 2: How to correct any misplaced points from the automated analysis of an AxoClamp file.

**Figure 18:**
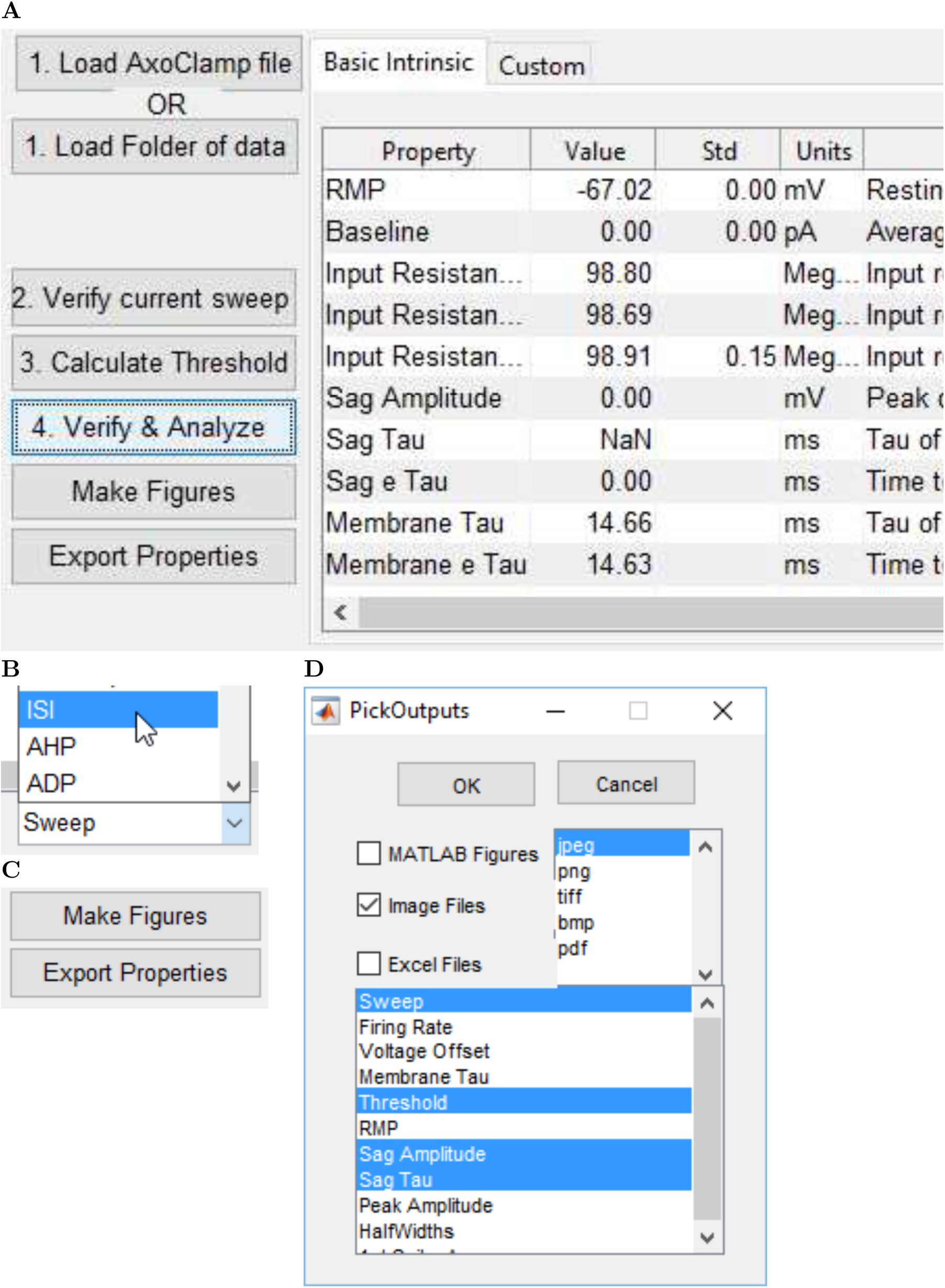
Tutorial 6, continued part 3: How to view and save figures generated from data in
an AxoClamp file.

**Figure 19:**
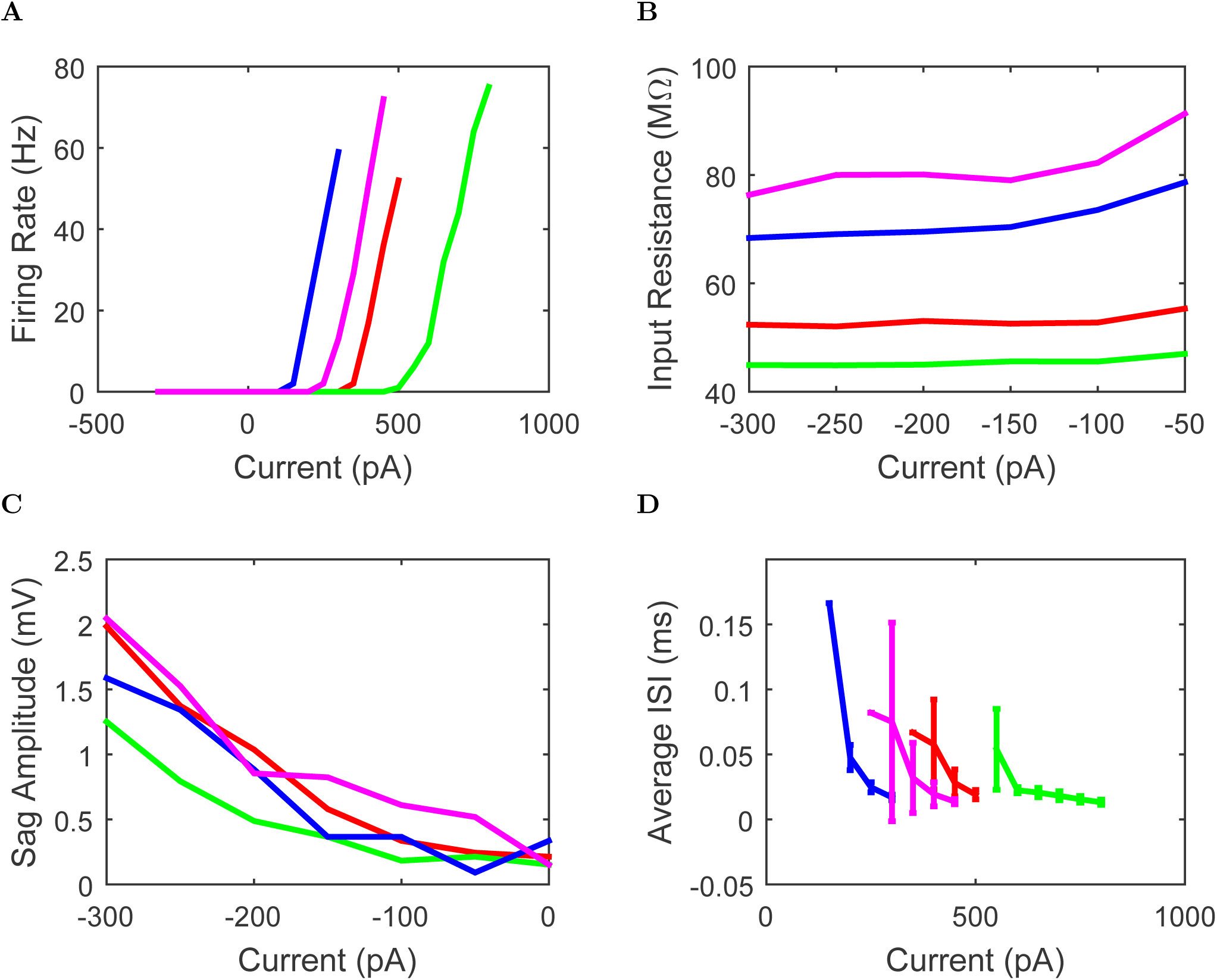
Example outputs from CellData automated analysis of a AxoClamp current injection sweep protocol. (A) Firing rates of the cells in response to current injections. (B) Input resistance of each cell as a function of injected current. (C) Sag amplitude in each cell as a function of current injection level. (D) Relation between interspike interval (ISI) and current injection.
17. Alternatively, click the ‘Export Properties’ button to export the table of electrophysiological properties as a tab-delimited text file, comma-separated values text file, or Microsoft Excel file.

**Figure 15:**
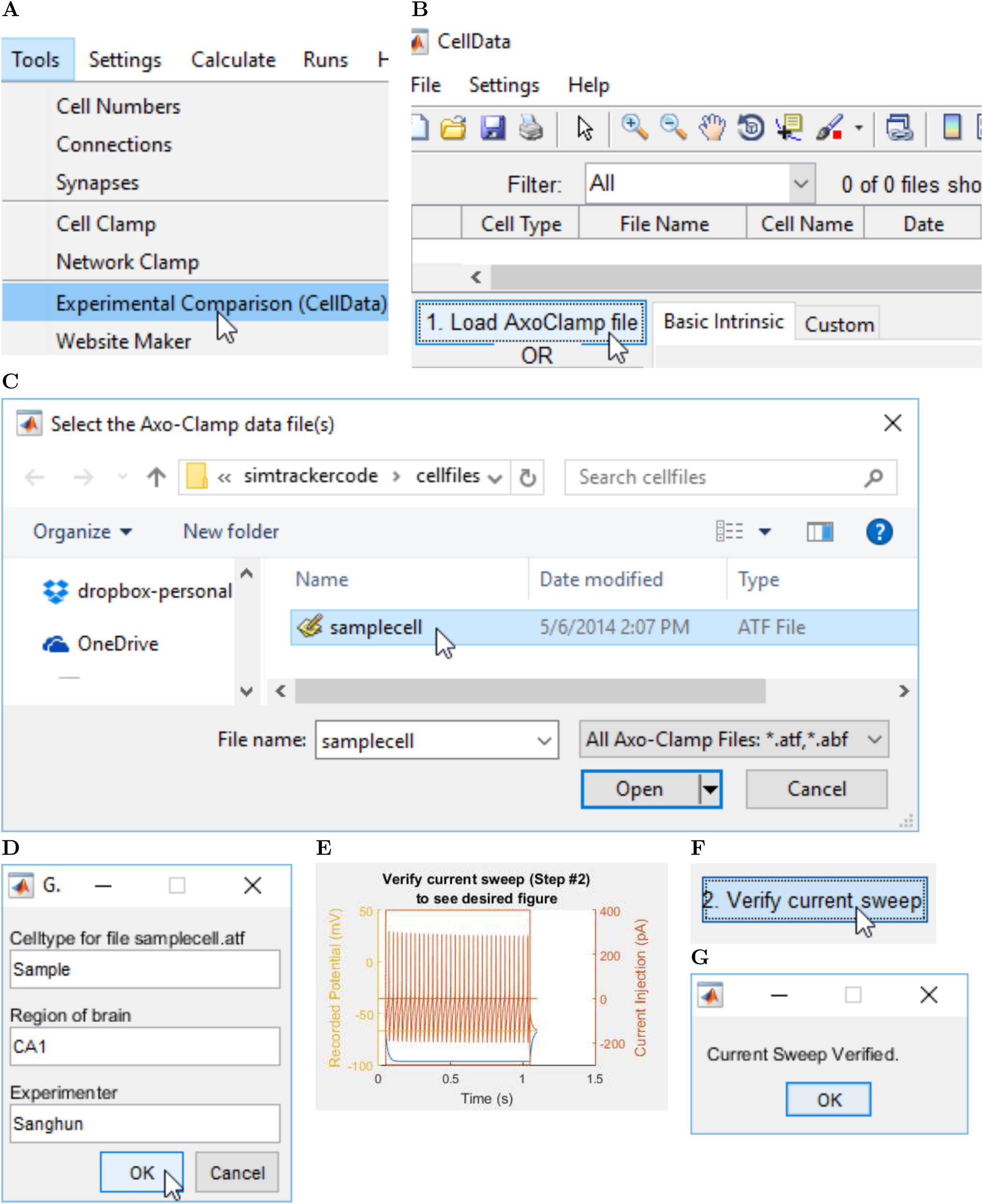
Passerine SINEs share a common ancestor and are mobilized by CR1-X.xs. (**A**) Maximum likelihood phylogeny of passerine SINE tails and avian CR1 subfamilies in Repbase (Jurka, et al. 2005) (GTRCAT model, 1,000 bootstrap replicates) suggests that TguSINE1 and PittSINE arose from the same CR1-X subfamily (CR1-X1_Pass) and share a common SINE ancestor. Note that the topology of the CR1 phylogeny is identical to that of previous studies (Suh, et al. 2012; Suh, Churakov, et al. 2015). (**B**) Comparison of the TguSINE1 landscape with landscapes of CR1 families (merged subfamilies from panel **A**) suggests temporal overlap of SINE and CR1-X activity in the zebra finch genome. RE landscapes were generated using the zebra finch assembly taeGut2 following methods detailed elsewhere (Suh, Churakov, et al. 2015).

#### Install SimTracker Manually

For users who do not wish to use an installation package, or who cannot find a package for their operating system, SimTracker and the associated applications can be installed manually on the computer by following the tutorial below.

### Tutorial 7: Install SimTracker and its associated software separately

Tutorial 7 will need to be completed on each computer that will run SimTracker. Some steps are only relevant on Windows computers and are marked ‘Windows Only’.

#### Software Requirements

To get the most out of SimTracker, users should download the following supporting software. For further assistance, please consult the detailed SimTracker user manual, available at http://mariannebezaire.com/downloads/SimTrackerInstructionManual.pdf or visit the websites of the individual software programs.

1. First, install the code versioning system Mercurial (https://www.mercurial-scm.org/) on the computer.
2. Next, install the NEURON simulation program (http://www.neuron.yale.edu/neuron/).
3. **Windows Only:** For Windows users, also install the terminal software Cygwin. Visit the Cygwin website at https://www.cygwin.com/ to download the installer and get detailed install instructions. Make sure to download the default packages as well as the packages below:
  - openssh
  - procps
  - rsync
  For extremely detailed instructions about downloading and configuring Cygwin to work with SimTracker, please consult the SimTracker user manual. However, SimTracker can guide users through most installation situations.

#### Download and install SimTracker

SimTracker is implemented in MATLAB, but potential users do not need to purchase a MATLAB. Non-MATLAB users can download the stand-alone, compiled version of SimTracker specific for their operating system type (Windows, Mac, or Linux). MATLAB users have the option of either downloading the compiled version or downloading the GUI scripts to use the SimTracker from within MATLAB. Both are available at the SimTracker website, http://mariannebezaire.com/simtracker/. Please consult the appropriate section below and then continue on to the next part of the tutorial.

##### Compiled SimTracker

We created a one-click installer for the stand-alone tool that installs both the SimTracker and the necessary MATLAB code libraries to run it. To use this installer:

1. Download the stand-alone installer file for the appropriate operating system from http://mariannebezaire.com/simtracker/.
2. Double-click or execute the file to launch the installer, then follow the prompts on the screen to install the MATLAB libraries and SimTracker. The installation may take a long time as there are many libraries to install, but it will alert the user when the installation has completed.

Note: On Windows, multiple prompts will appear warning the user about the downloaded installation file, due to its extension of ‘exe’. First, a warning will appear stating that the file MyPCAppIn-staller_web.exe is not commonly downloaded. For this particular file, the warning can be ignored. Do not discard the downloaded installation file. After launching the executable, another alert will appear stating that the file will not be run. Click the ‘More Info’ link for an option to run the file anyway. Finally, another dialog may appear asking if this executable file from an unknown publisher should be run. Click yes. Then, the installation should proceed normally.

Note: On Mac OS, users may need to change their security preferences to open the downloaded installer file. If a message appears that says ‘MyMacAppInstaller_web.app’ can’t be opened because it is from an unidentified developer, open the Mac settings, go to ‘Security and Privacy’, and look for a message that says ‘MyMacAppInstaller_web.app’ was blocked from opening because it is not from an identified developer. To the right of the message should be a button that says ‘Open Anyway’. Click this button (it may first be necessary to click the lock icon in the lower left corner of the security window prior to clicking the button). If the button does not work, the user may need to temporarily change the security setting to allow apps downloaded from ‘Anywhere’, then launch the installer, and then change the setting back to its previous, more secure option. When the installer launches, it will display another dialog requesting confirmation that the user wishes to download this application from the internet. Click the ‘Open’ button and the installation will proceed. The user will then be asked for their password to enable Java to set up the application on their computer. Remember to revert the security settings within Mac OS’s security dialog after installing SimTracker.

##### MATLAB Scripts for SimTracker

The first time users download the SimTracker code, they should download the compressed SimTracker code file, which contains all files necessary for the SimTracker to work. Users may incorporate future updates made to SimTracker by either downloading the compressed file again or obtain subsequent updates to the SimTracker code from the Mercurial repository located at https://bitbucket.org/mbezaire/simtrackercode. See the SimTracker user manual for detailed instructions on using Mercurial to keep SimTracker up to date. To obtain and run the SimTracker code:

1. Download the compressed file of the code from http://mariannebezaire.com/simtracker/.
2. Uncompress the downloaded file to the desired location by using either a WinZip-type utility or, at the command line, entering: `terminal-$ unzip SimTrackerCode.zip # (for zip files)` or `tar -xzf SimTrackerCode.tgz # (for tar files)`
3. Open MATLAB and change the current directory of MATLAB to the SimTracker directory
4. Type at the MATLAB command prompt type: `» SimTracker`

##### Configure SimTracker

Now launch SimTracker to choose a location for the model code and to configure SimTracker by reviewing the various settings of the tool.

1. After SimTracker launches, it will display a folder picker dialog. The user can navigate to an existing directory or create a new one using this dialog. Within the chosen directory, the user can store each model registered with SimTracker (in a separate subdirectory). SimTracker will also create other subdirectories within this directory as needed, for storing data backups, settings, and more.
2. Next, SimTracker’s General Settings dialog will appear, allowing the user to review the default configuration and make adjustments as necessary. In the window that appears, most settings can be left at their default. For now, only some settings under ‘Local System Commands’ must be updated as follows.
3. In the text field next to ‘Opens NEURON’, enter the path to the nrniv application on the local computer. If necessary, find this path by typing into the terminal: `terminal-$ which nrniv` On Windows, users must switch the slashes back to backslashes and replace the cygwin shortcut with the Windows Drive letter. For example, if Cygwin answers back /cygdrive/c/nrn73/bin/nrniv, then enter in the MATLAB setting C:\nrn73\bin\nrniv.
4. **Windows Only** For Cygwin users on Windows, also update the path to the Cygwin folder (the default installation is at C:\cygwin).
5. Below the text boxes, users should ensure the ‘Use Environmental Variables’ checkbox is selected unless they know what they are doing regarding environmental variables and wish not to have SimTracker fill in (overwrite) the values of PYTHONPATH, PYTHONHOME, NEURONHOME, and N for them.
6. Now, under the Jobs section, the user should enter an email address to which supercomputers can send notifications when jobs are complete.
7. Next, click the ‘Save’ button at top to close the General settings window.
8. Now, from SimTracker’s Settings menu, choose ‘Machines’.
9. In the list that appears, users should ensure that the name of their personal computer is listed. If it is not there, do the following:
  a. Click the ‘Add Line’ button
  b. In the new line, for the Machine name, enter the name of the computer. To find out the name, open the terminal (or Cygwin in windows) and enter the command: `terminal-$ hostname`
10. Leave the remaining fields in the line blank except for the last one, TopCmd. For TopCmd, enter “nrniv”.
11. Click the ‘Save’ button and then close the Machine dialog. The personal computer has now been added as an option. In addition to configuring SimTracker directly, it is necessary to create (or add to) a bash script in the user’s home directory, for use by the user’s terminal program. SimTracker will attempt to create a script if one does not already exist. If one already exists, a dialog box will appear after reviewing the general settings, instructing the user to add a line to the file to call an additional script for setting up SimTracker configuration. If the user prefers to follow these tasks manually, the following steps can be taken (substituting in the correct path for the user’s operating system and NEURON installation):
12. To find the home directory associated with the user’s account on their computer, open the computer’s terminal program (or Cygwin, on windows) and enter: `terminal-$ echo $HOME` and the terminal will display the home directory for that account
13. Within the home directory, create or open a file called “.bashrc” (users may name this file differently than .bashrc if necessary) using a code-friendly text editor (for users new to code editing, try Geany or Programmer’s Notepad 2, and make sure the application is set to use Unix line endings, not Windows line endings).
14. Within the file, add the following line(s), substituting in the paths shown below for the paths on the user’s particular computer (do not add the comments after the # sign, those are only to provide information for this tutorial) `export PATH= path/to/nrn73/bin:$PATH # the NEURON directory’s “bin” subdirectory` Users who unchecked the SimTracker general setting to ‘Use Environmental Variables’ during the tutorial above may also need to set (depending on their operating system) the four environmental variables below to the appropriate location for their computer:

~~~
export PYTHONHOME=/path/to/nrn73 # the NEURON or Python directory
export PYTHONPATH=/path/to/nrn73/lib # the “lib” subdirectory of PYTHONHOME
export N=/path/to/nrn73 # the NEURON directory
export NEURONHOME=/path/to/nrn73 # the NEURON directory
~~~
15. Save and close the file.

##### Access and run simulations on supercomputers

SimTracker can work with supercomputers to run large scale, parallel NEURON simulations such as the CA1 network model in (Bezaire et al., 2016). Below are several tutorials demonstrating the entire process of running a parallel model on a supercomputer. To follow along with these tutorials, the user must have access to a supercomputer. Here are three options, as stated in Bezaire (2014) at http://mariannebezaire.com/access-supercomputer/:

- Check if an affiliated company and institution has its own supercomputer or allows its affiliates access to another supercomputer
- Sign up for a Neuroscience Gateway (NSG) account; it is free, open to all, and takes care of the technical and administrative burdens that come with using a supercomputer so modelers can focus on their modeling work.
- Apply for an XSEDE startup grant. Award decisions are made relatively quickly and grant enough computing time to develop code into a full model and use the preliminary results to apply for a full XSEDE research grant.

When modelers run their code on the NSG, many of the technical details are already accounted for by the NSG. Therefore, we have provided a separate tutorial for running code on NSG that highlights the simplicity of the process, in addition to providing two tutorials for setting up NEURON on a supercomputer and then running a simulation on that computer.

### Tutorial 8: Run a model simulation on the NSG

Because the NSG portal takes care of NEURON code maintenance and all interactions with the batch queueing system, the process for executing a simulation via the NSG is slightly different.

1. First, select the simulation from the table at the top of the SimTracker, and then from the ‘Runs’ menu choose ‘Execute Run’ (same steps as for executing it on the local computer).
2. When the SimTracker displays a question box asking which NSG-linked supercomputer to use, push the button corresponding to the desired supercomputer.
3. A window will open with a zip file containing the configuration for the selected run. A dialog box with a list of job submission parameters will also open. Keep both the window and the dialog open until the simulation has been submitted.
4. Log onto the NSG portal at https://www.nsgportal.org/.
5. Navigate to the ‘Data’ section and click to upload new data. In the file navigator that appears, select the zip file corresponding to the one that is selected in the window that opened on the local computer. Upload the zip file to the NSG portal.
6. Navigate to the ‘Tasks’ section and create a new task. For the input data, select the zip file just uploaded. Then, for the values of the other job parameters, refer to the dialog box produced by the SimTracker. Set all the properties as indicated, and then click ‘Run Task’.
7. Users will want to occasionally check the status of their tasks in the NSG portal (and can also view the current output of the job in the ‘intermediate results’ section, stdout file).
8. When the job finishes, look at the stdout file in the output files section to ensure that the run completed successfully.
9. Then, download the large ‘.tar’ file of the results to the local computer.
10. Within the SimTracker, select the simulation and from the ‘Runs’ menu, choose ‘Upload Run’.
11. An empty results folder will appear with the same name as the simulation. Use the mouse to drag the downloaded tar file into that results folder (or use the command line to copy or move the file into that simulation-specific results directory).
12. Then, within the SimTracker, again select the simulation from the ‘Runs’ menu and again choose ‘Upload Run’. This time, because there is a ‘.tar’ file within the results folder, the file will be unpacked and all the regular results files will be transferred into the results folder, and then the unneeded files will be deleted. The simulation will now be uploaded into the SimTracker in the same way that the other simulations were uploaded.

### Tutorial 9: Prepare a supercomputer for use with NEURON and SimTracker

Users must first install NEURON on the supercomputer. It may be necessary to work with a system administrator to complete the installation. Technically-minded users may try the installation themselves by following the steps given in one of these forum posts:

- SDSC’s supercomputers (Comet and Trestles): http://www.neuron.yale.edu/phpbb/viewtopic.php?f=6…t=2792
- University of Texas’ TACC Stampede supercomputer: http://www.neuron.yale.edu/phpbb/viewtopic.php?f=6…t=2781

Next, set up SimTracker for use with the supercomputer. If using the compiled version of SimTracker, users are limited to using the supercomputers already programmed into SimTracker. However, if users are running SimTracker in MATLAB, new supercomputers can be added to the tool.

1. First, add the supercomputer to the Machines list in SimTracker by choosing ‘Machines’ from the ‘Settings’ menu.
2. In the window that appears, click the ‘Add line’ button and then scroll down to the new line in the top table.
3. In the first column, enter an easy-to-type nickname for the supercomputer with no punctuation or spaces.
4. In the second column, enter the web address of the supercomputer, either an IP address or the text address that would be used to access the supercomputer at the command line (such as stampede.tacc.utexas.edu)
5. In the third column, enter the username corresponding to the account on that supercomputer
6. Next, users must add the whole path to their ‘repos’ directory, which will house all of the models on that supercomputer
7. Optionally, in the fifth column users can add an allocation number (such as an XSEDE grant) to which the processor-hours used in their tasks will be counted against.
8. In the sixth column, type the number of cores (processors) included on a node of the supercomputer.
9. Next, enter the command used on that supercomputer to submit a job (job script) to the queue so that the simulation job can run when the requested number of processors become available. On many systems, the command is ‘sbatch’.
10. Leave the remaining columns of this line blank for now and click the ‘Save’ button.
11. Now, find the popup menu above the lower table and select the newly added machine from the list.
12. In this lower table, add job limits that must be obeyed when submitting the run for execution on a supercomputer (such as time limits or maximum number of nodes that can be requested). This lower table allows the user to list each queue available on this supercomputer and its limits. Be sure to type the name of the queue exactly as it is referenced on the supercomputer, and to add the hard time limits and maximum number of nodes if either of those apply. SimTracker will then look through the list of possible queues and request the first one it finds that fits the criteria estimated for the run. If no queues can accommodate the job configuration, SimTracker will display a message alerting the user of the problem rather than try to submit the job.
13. Then, click ‘Save’ again and close the Machines window. Users must next create a template for the style of job submission script that will need to accompany each simulation requested to run on the supercomputer.
14. In MATLAB, navigate to the ‘jobscripts’ directory within the SimTracker directory, and select and open one of the existing m-files within that directory (named with the convention ‘write-script_mysupercomputernickname’). Immediately save the m-file using the ‘Save As’ command to save it as a new file with the name ‘writescript_nickname’, where ‘nickname’ is replaced with whatever nickname the user gave the newly added supercomputer in the Machines list above.
15. Next, consult the user guide of the supercomputer to determine the necessary syntax and conventions of the job submission scripts that can be used on its system. Edit the newly saved m-file so that it produces a job script following the conventions of the documentation for the supercomputer. Users may wish to browse the other job script m-files already in the ‘jobscripts’ directory to see what variable names they reference to load information such as username, the queue name, allocation name, email preferences, number of processors, and other parameters.
16. Finally, save the changes to the new jobscript-creator m-file. The supercomputer is now ready to run a simulation.

### Tutorial 10: Run a model simulation on a supercomputer

This tutorial covers how to execute a simulation on a supercomputer.

1. From the machine list, users must choose a supercomputer to which they have access. Note that only supercomputers added by the user to the SimTracker using the Machine Settings dialog will be available in this list.
2. Next, specify the number of processors to use for the job. This will depend on how much memory the model needs, how much memory each supercomputer node can provide, the number of processors per node on the supercomputer, and the time requirement for the model. The most basic strategy for determining what number to use would be to run a scaled down network for a short amount of time to gauge the memory usage and time requirements of the model. For further guidance on setting this number, see the SimTracker Instruction Manual.
3. Then, considering of how much time will be needed to run the model with the given number of processors, pad the estimate and enter the number in hours for the JobHours parameter. For example, if confident the simulation should take 4 hours to run, consider entering 5 or 6 hours in the JobHours field. Parts of hours should be entered as decimals (i.e., 30 minutes would be entered as 0.5, 90 minutes would be entered as 1.5).
4. Separately, enter a number of hours for the EstWriteTime field. This parameter sets the number of hours that will be reserved for writing from the user’s JobHours parameter. For example, if the user entered 5 hours for JobHours and 1 hour for EstWriteTime, the simulation will end early if it is in danger of taking more than 4 hours, allowing a full 1 hour to write results to disk. This helps ensure that the simulation results are not completely lost if the model runs slower than anticipated and is in danger of being terminated due to the hard time limit coded in the job submission script.
5. Depending on personal preference and also the behavior of the supercomputer, users may wish to disable the model code’s system calls. While the system calls coded into the model help to ensure the proper documentation of each simulation and also help streamline the organization of the simulation results, for some users they may produce unacceptable side effects. If desired, they can be disabled by setting the CatFlag parameter to 0.
6. Then click the ‘Execute’ button. Users may be prompted to enter the supercomputer account password either at the MATLAB command line or into a dialog box that pops up. Users may receive multiple requests for their password (due to SimTracker issuing multiple commands to the super-computer) if they have not set up ssh keys or GSI for their supercomputer account.
7. If the simulation has been submitted to a supercomputer not using the NSG, a dialog box will appear after submission alerting the user that the run has been submitted to the supercomputer. Best practice is to log into the supercomputer after receiving that message and check its queue periodically to ensure the simulation has been successfully executed.
8. Users may want to monitor the queue on the supercomputer to see when the simulation begins and finishes. They may also set up the job scripts to email themselves whenever the run finishes (see previous tutorial for assistance on creating and modifying job submission scripts). Once the run finishes or disappears from the queue, view its output log file to ensure the run executed successfully.
9. After receiving a confirmation email or seeing that the run has completed, log into the supercomputer to check that the run succeeded.
10. Once logged in the supercomputer, find the repository and view the last few lines of the output script produced by the supercomputer for the run, using a command such as tail -n 20 ./job-scripts/MyRunName.*o. The summary lines are always printed at the end of a run.
11. Then, in SimTracker, select the run and from the Runs menu, choose ‘Upload Run’. It may take a while to transfer all the files from the supercomputer to the local computer. The program will compress the results file on the supercomputer, transfer the compressed file over, and then unpack all the files out of the compressed package on the personal computer. For large simulations with hundreds of thousands of cells, lasting several seconds, and especially if the user chose to record the voltages from several hundred or thousand cells, the upload process may take 30 minutes to an hour.
12. After the results have uploaded, a dialog box will appear stating that the results are uploaded. Users may then wish to look at the time spent on the run, the time spent by each processor for each major section of the code, the load balance, and the variation in times for different processors to complete different code sections. Large variations or low load balances (<.75) may indicate that the code is not well parallelized.

For additional tips on increasing efficiency when working with supercomputers, such as configuring auto-login for a supercomputer and checking the status of a simulation on the supercomputer, see the SimTracker users’ manual available at http://mariannebezaire.com/downloads/SimTrackerInstructionManual.pdf. In addition, please see the instruction manual for troubleshooting tips about supercomputers or specific to SimTracker. For more general discussion of parallelizing NEURON models, please see (Hines and Carnevale, 2008).

#### Model Design Characterization

Here, we demonstrate how to add cell types and alter the network constituents or connectivity. The following tutorials also highlight additional tools and features of SimTracker. These tutorials work best with a larger network model rather than the ringdemo network. Feel free to start a new repository using the ca1 model code by following Tutorial 3 and selecting ‘ca1’ rather than ‘ringdemo’ when adding the new repository to SimTracker. Then proceed with the following tutorials.

### Tutorial 11: Add model cell types and specify model cell numbers and connections

This tutorial will illustrate the necessary steps to add another interneuron type to the ca1 model network, such as perforant path-associated cells (PPA cells). First, we would need to create a new model cell template for this cell type, and then we would need to create some datasets containing information about how to incorporate that cell type into the model network. We will walk through both of these goals in the following tutorial.

The tutorial will begin by copying an existing, working model cell template. It is a good idea to create the model cell template by adapting it from an existing file. Users can make incremental changes to a copy of the working template to ensure that they maintain the standardization and functionality of the template and also to ensure the ability to check that the template still works after each change.

1. First make a copy of the Schaffer Collateral-Associated (S.C.-A) cell template, as the S.C.-A. cell type is quite similar to the PPA cell type. Within the ‘cells’ subdirectory of the model repository, open the ‘class_scacell.hoc’ file and choose ‘Save As’ and then type ‘class_ppacell.hoc’. Now the new cell template is ready to be edited and tuned to become a model PPA cell.
2. Next, rename the template within the file. The first and last code lines of the file refer to the template name, which should be ‘ppacell’ instead of ‘scacell’. To work most efficiently, use find and replace to change all references to ‘scacell’ to ‘ppacell’ in the document.
3. Next, alter the morphology and electrophysiological properties defined in the file as we see fit. Later on, there will be an opportunity to test and further tune the cell using the CellClamp tool to ensure that it behaves as a physiological PPA cell.
4. Finally, save the file and then return to the SimTracker tool to add the cell to the network model. Now that the model PPA cell template has been created and added to the ‘cells’ directory, it is time to create the new datasets that will include this cell type in the network.
5. From the SimTracker ‘Tools’ menu, choose the ‘Cell Numbers’ option.
6. A dialog box will open, allowing the user to select an existing cell numbers set or to create a new set that specifies which cell types to include in the model and how many of each should be included. To create a new set based on the existing set #101, choose 101 from the top menu to load that cell numbers dataset.
7. Before making any changes, click the ‘Save New…’ button and add a comment to the box that appears, stating ‘#101 with ppacell added.’ This will save a new dataset that can be edited.
8. Now, from the celltype menu, select ‘ppacell’ on both sides and then click ‘Add’.
9. A new line will be added with an entry for ‘ppacell’, Enter the remaining information on that line, including the number of ppacells in the full size network and the layer in which they are found (stratum lacunosum-moleculare is layer 3, since stratum oriens is layer 0).
10. Next, click the ‘Save’ button. A dialog box will appear stating that a new axonal distribution file has been created for the ppacell. To specify the axonal extent and distribution of boutons of the ppacell, modify its axondist file from within the axondists directory of the cells directory in the repository. Further information about this file and its parameters is included in the User Manual.
11. Now, close the Cell Numbers dialog box.
12. Next, from the SimTracker ‘Tools’ menu, choose ‘Connections’. A matrix will appear. From the top left menu, select ‘430’ to load the basic dataset from which we will derive our new one. Then, from the top right menu, select the number corresponding to the new cell numbers dataset just saved. The matrix will now expand to include ppacells as well. Down the left side of the matrix (the row labels) are given the presynaptic cells, while the top row (the column labels) gives the postsynaptic cells. At the intersection of each presynaptic and postsynaptic cell, add the average number of connections from all cells of that presynaptic type onto one postsynaptic cell of that type (aka, the convergence).
13. After filling out all the connection information for connections to or from ppacells (convergence numbers, synapses per connection, and synaptic weights), save the connection tool and close it.
14. Finally, from the ‘Tools’ menu choose ‘Synapses.’ A table will appear. From the top left menu, select the number ‘120’ to load a baseline synapse dataset.
15. Then, from the top right menu, select the number corresponding to the recently created cell numbers dataset that contains the PPA cells.
16. Now, notice the pop-up menu at the top middle to specify a post-synaptic cell. Select the ppacell from this menu and then add potential presynaptic cells to the list to specify their specific kinetics. Save the dataset by clicking the ‘Save’ button.
17. Now, switch the top middle menu to each cell that can receive input from the PPA cell in turn. Add the synaptic entry for the ppacell connection. Then, before switching to a different post-synaptic cell, again save the current table. After completing the definitions for all postsynaptic celltypes, then save the table again and push the button to close. One of each type of the parameter dataset has been created to specifying the network and its connections: cell numbers, connections, and synapse kinetics. These datasets resemble those used in the control CA1 network simulation except that they also include PPA cells. Now, users can launch the CellClamp tool, set the dataset menus in it to the ones just created, and test both the intrinsic cell properties of the PPA cell and the connections to and from PPA cells (using the paired recording feature). The next tutorial covers the use of the CellClamp tool.

### Tutorial 12: Characterize model network components with CellClamp

Here, we walk through how to characterize basic components of the model using the Cell Clamp tool. This tool can characterize ion channel dynamics, single cell intrinsic properties, and synaptic connections in common experimental terms. This tutorial shows each of these characterizations using the CA1 network.

1. From the SimTracker Tools menu, choose ‘Cell Clamp’. The Cell Clamp tool will open.
2. Within Cell Clamp, first, scroll through the table of ion channels to find the entry for Nav, or the voltage-gated sodium channel. Fill out the entries for this row in the table, entering a Gmax (maximum conductance density) value of 0.001 (micro Siemens per square centimeter) and an ENa (reversal potential) value of +55 (mV). The variable used to store the reversal potential for this channel, ena, has already been populated in the table.
3. Next, click the checkboxes to the right of the ion channel table, for ‘IV Curve’ and ‘Act./Inact.’. Then, click the ‘Conductance’ radio button below them so that the results will display in terms of ion channel conductance rather than current.
4. Then, click the ‘Get Results’ button below. A dialog box will appear, asking the user to add comments for this particular characterization run. Enter ‘Characterize Nav channel in conductance terms’ and then click ‘OK’. It will take a little while for the characterization simulations to complete before the result figures appear. Because the Nav row in the table was filled out, the CellClamp will run an activation/inactivation curve and a current/voltage relation for the Nav channel and display the resulting graphs in terms of channel conductance.
5. Next, prepare CellClamp to run a simulation at the single cell level rather than the macroscopic ion channel level. Uncheck the ‘IV Curve’ and ‘Act./Inact.’ checkboxes so that these protocols will not run again, and instead check the ‘Current Clamp’ checkbox to the right of the list of cells (in the middle of the tool).
6. Check the popup menu above the list of cells and set it to the ‘cellnumbers_100’ dataset. All the cell types included in that dataset will populate the list of cells below.
7. From the list of cells, click to select the ‘pyramidalcell’ cell type.
8. Set the current clamp protocol to be applied to the selected cells. The top row of numbers gives the time before the pulse is applied, then the length of the pulse, followed by the time after the pulse has finished that the cell behavior continues to be recorded. The row below gives all the current injection (or voltage clamp) levels that will be used. Note that MATLAB’s vector syntax can be used to specify which currents to apply. For example, rather than listing −0.300, −0.250, −0.200, −0.150 … all the way up to +0.500, users can specify that every current injection level between -.300 nA and +.500 nA should be tested, increasing in steps of 0.050 nA, with the following syntax: [-0.300:0.050:0.500]. Users can also add additional values to use before or after that syntax within the brackets: [−0.500 −0.300:0.050:0.500 0.510 0.540 0.580 0.600]. For this tutorial, leave the row of times alone but change the current list to say: [−0.3:0.05:0.5].
9. Now click ‘Get Results’ and enter into the comments dialog that appears ‘Single cell characterization’. After a few moments, several figures will appear, characterizing the cell’s behavior. Note: For even more in-depth characterization, the results of this CellClamp protocol can be loaded into the CellData tool to compare the model cell’s properties directly with those of experimental cells.
10. Next, prepare the CellClamp tool to run a paired recording rather than a single cell recording by unchecking the ‘Current Clamp’ checkbox in the single cell section.
11. In the lower left area of CellClamp, set the popup menu to ‘syndata_120’ to populate the synapse list below with all synapses listed in that dataset.
12. From the synapse list, select the pvbasketcell -> pyramidalcell entry to characterize connections from PV+ basket cells to pyramidal cells.
13. Click the current clamp checkbox to apply a current clamp to the postsynaptic cell in the pair (the pyramidal cell), and to the right of the checkbox, enter ‘0’ (nA) into the current injection box so that the cell’s response to the incoming synaptic activity will be recorded relative to its baseline resting potential.
14. Above the current injection amount, the row of boxes indicated relevant times for the paired recording protocol: 15 ms after the recording starts, the presynaptic cell will spike, triggering the synaptic activity in the postsynaptic cell after a short delay due to axonal conduction. The next entry, 100 ms, shows the length of time to record the postsynaptic cell’s response after the presynaptic spike. For most current clamp of synapse types, 100 ms is adequate; only for slow synapses such as those with GABAB would the recording window need to be longer in a current clamp paired recording.
15. Finally, click the ‘Get Results’ button and for the comment dialog, enter ‘Synaptic recording’ and then click ‘OK’. The CellClamp will now perform 10 paired recordings of the synaptic connection using the protocol specified, and then will average the results. It will then display a graph of the individual recordings and the averaged recording, along with the time constants and amplitude of the synaptic connection computed from the averaged trace.

### Tutorial 13: Network Clamp a cell using the NetworkClamp tool

Next, use the Network Clamp tool to employ the Network Clamp technique (Bezaire et al., 2016) on the model. The Network Clamp is most useful for a large network, especially one with many inputs to each cell. The Network Clamp tool can be used for investigating how various inputs affect individual cells and for studying network dynamics without requiring a supercomputer. For this example, we will study how a particular cell’s behavior changes as a function of the strength of the incoming synapse. This will gives us an idea of how to alter our connectivity strength in the full model without having to run a lot of full-network simulations to find the proper synapse strength. We will run this tutorial from the ca1 repository, so please ensure that SimTracker currently has the ca1 repository open before proceeding. The first few steps of the tutorial will download the results and configuration (Bezaire et al., 2015) from the control, full scale CA1 network run used in Bezaire et al. (2016), so that they can be used as the basis for a network clamp run in the rest of the tutorial.

1. First, create a free account on the CRCNS website at http://crcns.org/data-sets/sim/download.
2. Then, visit the entry for the simulations results of Bezaire et al. (2016) at http://crcns.org/data-sets/sim/sim-1/about-sim-1, enter the name and password, and download the results file for ca1_centerlfp_long_exc_065_01.
3. Then, from within SimTracker, choose File > Import and in the file picker that appears, select the newly downloaded ca1_centerlfp_long_exc_065_01 file to import it. The simulation run should then appear in the SimTracker runs table.
4. From the SimTracker Tools menu, choose ‘Network Clamp’ and the Network Clamp tool will open.
5. Within the Network Clamp tool, at the top left area in part “1. Inputs”, select the ‘Simulation Results’ radio button.
6. Then, from the popup menu next to it, choose the network simulation to use as the baseline: ‘ca1_centerlfp_long_exc_065_01’ and change the duration from 4000 to 100 (ms).
7. Next, find the popup menu for ‘Use inputs for’ and choose ‘pyramidalcell (21310 – 332809)’, and for gid number, enter 21310.
8. For part “2. Electrophysiology”, choose the electrophysiology of ‘poolosyncell’, which is the technical cell template used in the model CA1 network, derived from the pyramidal cell model published by Poolos et al. (2002).
9. Finally, for part “3. Connections”, choose to use the connections and synapses from the corresponding network simulation by clicking the ‘From Run’ radio button for the Connections line as well as for the Synapse kinetics line. Then verify that the ‘Run’ menu below says ‘ca1_centerlfp_long_exc_065_01’, as the simulation run from which to take the connections and synapse kinetics.
10. At the bottom left area, enter a description for this Network Clamp run, such as ‘Running baseline netclamp for pyramidal cell’, click the ‘Table View’ radio button for ‘Results’ and then click the ‘Execute’ button.
11. After the run completes, the intracellular membrane potential recording for the cell will appear on the graph on the right side, as well as histograms of each input type received by the cell throughout the simulation. In the table at the top right, a summary of the run will appear. Users can toggle whether the design or the results of the network clamp run are shown using the radio buttons at the bottom left of the NetClamp tool. Next, try altering the inputs to this cell and observing the effect. This tutorial will arbitrarily focus on investigating the effect of distal feedback inhibition, which is mediated by neurogliaform cell inputs on the pyramidal cell.
12. In part “3. Connections” on the left side of the NetworkClamp, switch the connections radio button to ‘Custom Table’. Then in the table below, find the row for input cell “ngfcell” (for neurogliaform cell) and the column for ‘Wgt (uS)’. Change the value from 1.4500e-04 to 1.4500e-05, decreasing by 90% the incoming weight of inhibition from neurogliaform cells to the network-clamped pyramidal cell.
13. In the description field at the bottom left of the tool, enter a new description of ‘90% reduction of neurogliaform cell input’ and then click ‘Execute’ button. The simulation will run again with all the same times of spike inputs to the pyramidal cell, and the incoming synaptic weights and kinetics will remain the same for all inputs except for neurogliaform cell inputs. The neurogliaform cell inputs will occur at the same time, with the same kinetics, but have a synaptic amplitude that is only 10% of the baseline condition.
14. After a few minutes, a new record will appear in the upper right table and a new trace will appear on top of the old one in the graph just below it. The input histograms will update to reflect the inputs from this new run.
15. Various run results can be hidden or displayed by clicking the checkbox next to the corresponding description in the results table. The color and dash pattern of each results line can also be customized by adjusting the settings in columns “Col” and “Line” of the results table. Currently, the input histograms always display the input spike times from the most recent network clamp simulation run.

#### Model Component Tuning

After performing a characterization of a model cell using CellClamp or Network Clamp, users may discover that certain model components do not adequately fit their experimental counterparts. In this situation, SimTracker can aid in the tuning the cell. The MRFHelper, currently available from within the CellClamp tool within SimTracker, can be used to prepare a tuning protocol that uses NEURON’s Multiple Run Fitter (MRF). MRFHelper allows users to specify an experimental recording to fit and the parameters of the model that can be varied, and then it creates a session file that can be opened in NEURON, launching a pre-configured MRF tool which can be used to optimize the model cell as explained in the following tutorial. The advantage of MRFHelper is that it saves users from having to do the manual preparation involved with setting up a new MRF session in NEURON, and provides a gentle introduction to the capabilities of MRF.

### Tutorial 14: Use CellClamp to configure a MRF session in NEURON

The MRFHelper is currently accessible within CellClamp.

1. From the “Experimental Data” menu of CellClamp, choose “Prepare MRF”. A form will appear that allows the user to specify the experimental data to fit, the model cell to tune, and the parameters to vary in search of a good match between model and experiment.
2. In the “Biological Cell” menu at top, select the experimental cell to fit. (The cells listed in this picker menu come from the experimental cell data uploaded to the CellData tool.)
3. Next, pick the property or data to fit. Currently, the MRFHelper only tunes model cells to membrane potential traces recorded during a single current injection level. However, this functionality will be updated in the future, and users who are comfortable with NEURON’s MRF tool can configure MRF sessions to tune cell properties instead of raw data.
4. Add a current base if applicable, which will be applied to the cell throughout the entire simulation, in addition to the current injection level selected above.
5. Select the model cell to be tuned from the “Model cell” picker.
6. Click “Generate” to set up the MRF session. All the necessary files will be generated within the model repository/cellclamp_results/MRF directory. In that directory will be the following files:
  - a file of the biological data to fit: biodata_[cellname].dat
  - a file defining the model cell: modelcell.hoc
  - a file to initialize the simulation environment: MRFinit.hoc
  - a file of instructions for starting the MRF session: MRFinstructions.txt
  - a file to define the NEURON session: myMRFsession.ses
  - two supporting files used to define the session: myMRFsession.ses.fd1 and myMRFsession.ses.ft1
7. Follow the instructions in the MRFinstructions.txt to start the MRF session. Briefly, the user may open a terminal program, change the directory to the model repository, and then enter the command:

~~~
terminal-$ nrniv cellclamp_results/MRF/myMRFsession.ses
~~~
8. NEURON will open and load the MRF session, including several windows. First look over the windows to ensure the parameters and settings look ok.
9. Finally, to start the tuning process, find the window called MulRunFitter Optimizer and click the “Optimize” button.
10. To learn more about how the MRF tool works and how to use it, please see the tutorial available at http://www.neuron.yale.edu/neuron/static/docs/optimiz/model/set_up_runfitness.html

#### Publishing Model Results

Finally, we can make broadly available the completed characterizations and simulations of our network. The figures and tables produced from the SimTracker output can be used in publications, but the structure of our results also lends itself to being displayed on the internet. The next tutorial will use the WebsiteMaker to populate the results of our characterizations into a website template that is part of the SimTracker package. The website code generated can then be put directly online to help describe a model that a user has created.

### Tutorial 15: Create an interactive website to browse the network model

After running characterization protocols on all model cell types, ion channel types, and possible connections, the user may wish to show how the behavior of each model component or compare the model component with experimental observations. We have created and provided a website template that can be populated with results from the model characterizations performed by the user within the CellClamp tool. The resulting website is intuitive, well-organized, and provides a thorough overview of the model components and their fit to the experimental data where applicable (Figure 20).

Note: the Website Maker tool is meant to be used after the user has run all desired characterization protocols in CellClamp, as it will require the user to choose from the completed characterizations for posting on the website.

1. From the Tools menu of SimTracker, choose ‘Website Maker’.
2. A separate window will appear, listing simulations and CellClamp characterizations to be selected for inclusion on the website.
3. First, choose a baseline simulation from the drop down menu in the top right corner. The datasets from this simulation will be used to populate the tables on the website regarding cell types and numbers, incoming and outgoing connectivity, and weights and kinetics of synapses.
4. Next, if there is a publication associated with the model or website, enter its citation and URL link in the corresponding fields. Now the particular simulations and characterizations can be chosen from the lists along the bottom half of the window. Multiple selections are allowed in each list. As this tool can also be used to update an existing website, checkboxes are included to allow the user to only update a portion of the website (checkboxes above each list). Users also have the option to only update a subset of data for a portion of the website (default), or to completely replace the portion with the selected cells (by selecting the ‘only’ button below each list). For example, a user may have a current website that includes information about many cell types, but then run the Website Maker and only select the pyramidalcell type from the cell list. If the user runs the Website Maker that way, the pyramidalcell part of the website will be updated while the other cells’ data will remain untouched and intact. However, if the user also selects the ‘Only’ checkbox below, all cell data will be removed from the website and only the updated pyramidalcell data will remain.
5. In the bottom left list, select all simulations whose results should be included in the website and select the box above the list.
6. In the next column, select the cell characterizations to use on the website. The cells chosen should have had their CellClamp results imported into CellData and analyzed using the standard scripts for analyzing biological cells. Also select the box above the list.
7. Next, select the paired recording simulations run in CellClamp at the common, somewhat physiological condition. Including this characterization on the website allows different synapses to be compared to each other under similar conditions, unlike the results from varying biological experimental protocols. Again select the box above the list.
8. In the second from right column, select the paired recordings that were generated under conditions corresponding to various experimental protocols. This allows relevant comparisons of model and observed, biological cell behavior. Also select the box above the list.
9. Finally, in the right column, select the ion channel protocols to publish on the website and select the button above the list.
10. Now, click the Create button to create a website based on the simulation and characterization selections.
11. When the Website Maker is finished, a dialog box will appear giving the location of the website code. This code must now be uploaded to the server (or website host account) using an FTP program (such as WinSCP for Windows or Fetch for Mac). The main page of the site, which will be the end part of the link to get to the site, is called ‘mymodel.html’ by default, but it can be renamed if the user desires. The website will look similar to the existing one for the ca1 network model, pictured in Figure 20

